# Mammalian promoters are characterised by low occupancy and high turnover of RNA polymerase II

**DOI:** 10.1101/2024.09.23.614464

**Authors:** Kasit Chatsirisupachai, Christina J.I. Moene, Rozemarijn Kleinendorst, Elisa Kreibich, Nacho Molina, Arnaud Krebs

## Abstract

The general transcription machinery and its occupancy at promoters are highly conserved across metazoans. This contrasts with the kinetics of mRNA production that considerably differ between model species such as *Drosophila* and mouse. The molecular basis for these kinetic differences is currently unknown. Here, we used Single Molecule Footprinting to measure RNA Polymerase II (Pol II) occupancy, the fraction of DNA molecules bound, at promoters in mouse and *Drosophila* cell lines. Single molecule data reveals that Pol II occupancy is on average 3-5 times more frequent at transcriptionally active *Drosophila* promoters than active mouse promoters. Kinetic modelling of the occupancy states suggests that these differences in Pol II occupancy are determined by the ratio between the transcription initiation and Pol II turnover rates. We used chemical perturbation of transcription initiation to determine Pol II turnover rate in both species. Integration of these data into the model shows that infrequent Pol II occupancy in mammals is explained by the combination of high Pol II turnover and low transcription initiation rates.

## Introduction

Precise control of the levels and the timing of gene expression is required for the successful development and homeostasis of organisms. The production of a transcript by RNA Polymerase II (Pol II) is a sequential process started by the opening of the chromatin at promoters and the binding of general transcription factors (GTFs), which form the pre-initiation complex (PIC) and recruit Pol II onto the DNA. Upon initiation, Pol II transcribes for a few nucleotides and pauses to enable mRNA capping. Pol II is then either released into elongation to produce a full-length transcript, or terminated by the integrator complex (1, 2). Transcription is a discontinuous process that occurs in bursts. Active genes typically switch between an active state where transcription occurs, and inactive states during which the gene is silent (3). Each gene has a characteristic bursting pattern with a given frequency and intensity that, together with the mRNA decay rate, defines the steady state levels of mRNA in cells (4–6). Although gene bursting is conserved across eukaryotes, it happens on strikingly distinct timescales in different organisms. For instance, in *Drosophila* embryos the duration of the active and inactive state are in the same order of magnitude at the minute scale (7–10), while mammalian genes typically have long inactive states, ranging from tens of minutes to hours, interspersed with short active states (11–14). The molecular basis for these wide differences in transcription kinetics is yet unknown (15, 16).

A possible explanation could be the divergence in the molecular mechanisms underlying transcription. Yet, biochemical and structural data argue for a high degree of conservation of the protein complexes regulating the various steps of transcription across metazoans. For instance, the protein complexes controlling transcription initiation, namely the PIC and mediator complex, are highly conserved from yeast to human (17–22). In addition, the factors that control entry into elongation through polymerase pausing, such as the DRB sensitivity inducing factor (DSIF), the negative elongation factor (NELF), and the positive transcription elongation factor b (P-TEFb), as well as the Integrator complex that mediates early transcription termination, are conserved across metazoans (23–26).

An alternative hypothesis is that the same set of molecular complexes are used, yet with different kinetics. The long periods of transcriptional inactivity characterising mammalian genes suggest that, at any given time, promoters will be active in only a small fraction of cells within a population. This could be explained by the fact that promoters are experiencing initiation less frequently and consequently have lower occupancy by GTFs and Pol II. Another explanation could be that genes have similar rates of initiation, but that they are more frequently subject to non-productive transcription through increased rates of Pol II pausing or premature transcription termination. Pol II was found to accumulate shortly downstream of the of active genes transcription start site (TSS) in both *Drosophila* and mouse (23, 27, 28), suggesting no striking differences in the distribution of Pol II at sites of initiation, pausing or entry into elongation. Thus, there is an unresolved paradox between the large difference in the kinetics of transcription and the similarity of the molecular mechanisms supporting this process. It illustrates the challenges in connecting data from dynamic measurements of transcription kinetics with the static view of the genome occupancy of transcription regulators obtained by genomics.

Genome-scale studies of GTF and Pol II occupancy have been mostly based on bulk genomics assays such as chromatin immunoprecipitation sequencing (ChIP-seq) or precision run-on sequencing (PRO-seq). These assays can reveal the relative occupancy of factors across the genome at nearly base pair resolution (29–32) and they have been widely used to compare the relative abundance of Pol II at different promoters or between different experimental conditions. However, a limitation is that these assays are based on the sequencing of DNA that is experimentally enriched, representing an average occupancy over millions of cells. In addition, the enrichment of a protein such as Pol II at a promoter cannot be translated to the proportion of cells in which this binding occurs (Figure 1A, left). This makes it challenging to use these methods to compare the absolute promoter-proximal protein binding levels between species with distinct genomes, such as invertebrates and mammals.

**Fig. 1.**
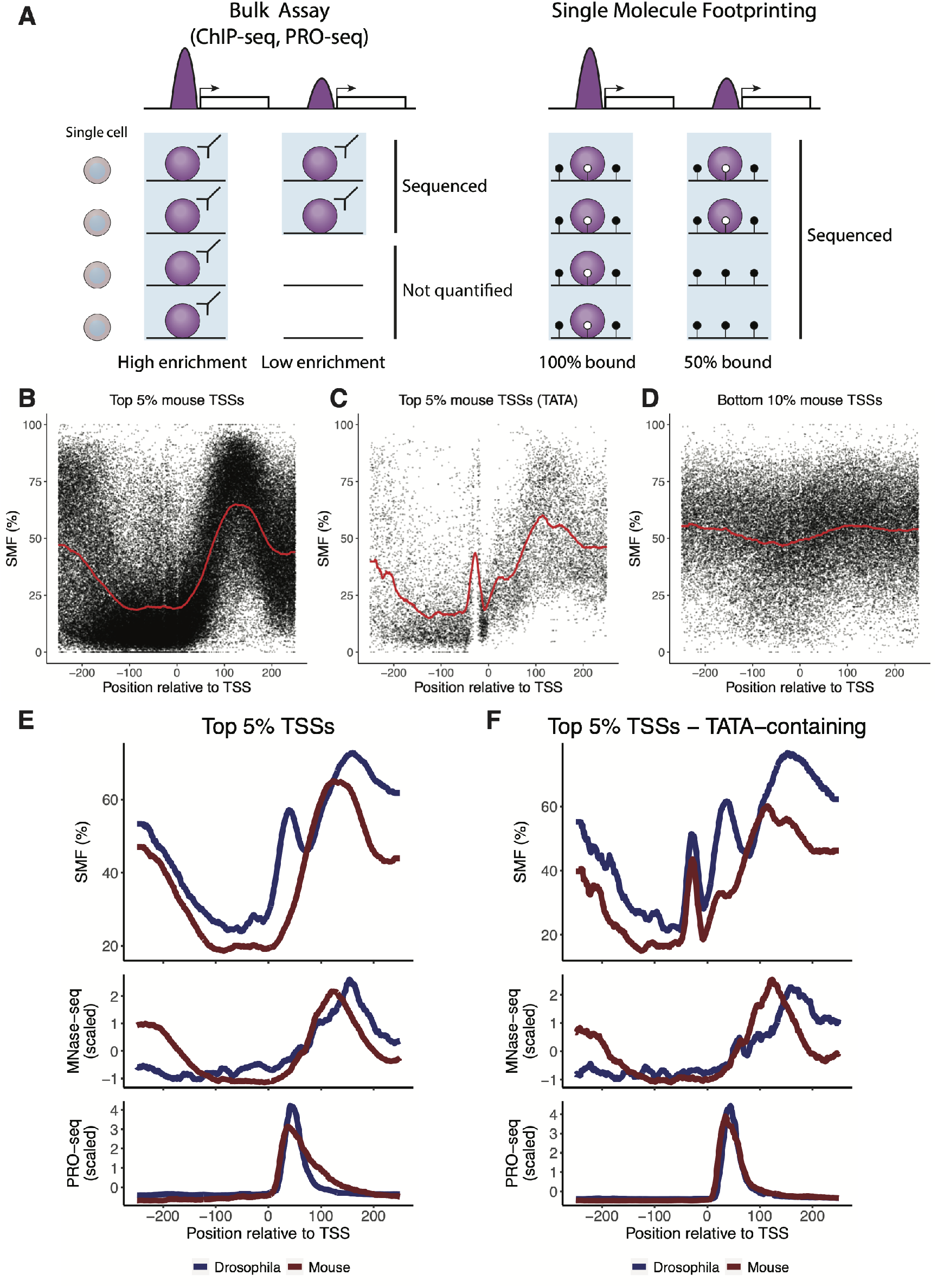
SMF reveals the difference in average Pol II occupancy at mouse TKO mESC and *Drosophila* S2 cell promoters. **(A)** Difference in data modality between bulk genomic assays and the single molecule assay. While methods such as ChIP-seq or PRO-seq report an enrichment, SMF allows absolute quantification of binding frequencies in the cell population. The light blue box indicates the DNA fraction that is sequenced with an assay. **(B-D)** Composite profile of SMF signal (1 - methylation [%]) at the **(B)** active mouse TSSs (Top 5% Pol II ChIP-seq, n = 1,316 promoters), **(C)** active TATA-box containing mouse TSSs (Top 5% Pol II ChIP-seq with a TATA-box, n = 151 promoters), and **(D)** inactive mouse TSSs (Bottom 10% Pol II ChIP-seq, n = 2,420 promoters). Each dot represents an individual cytosine. The red line indicates the smoothed average signal over 20 bp. **(E-F)** Comparison of the average SMF, MNase-seq, and PRO-seq levels between mouse (red lines) and *Drosophila* (blue lines) of **(E)** top 5% TSSs (n = 1,316 promoters for mouse; n = 931 promoters for *Drosophila*) and **(F)** top 5% TATA-containing TSSs (n = 151 promoters for mouse; n = 152 promoters for *Drosophila*). For MNase-seq and PRO-seq, reads at each position relative to the TSS [-250:250] were normalised to reads per millions (RPM). The RPM at each position relative to the TSS were averaged across top 5% and top 5% TATA-containing TSSs. The average RPMs were then normalised to the Z-score to enable a comparison between mouse and *Drosophila* in the same plots.

Single Molecule Footprinting (SMF) quantifies protein-DNA contacts at single molecule resolution across the genome (33, 34). The single molecule resolution of SMF enables the direct quantification of the frequency of promoter occupancy by various DNA binding proteins including transcription factors (TFs), GTFs, and Pol II (Figure 1A, right) (34, 35). We showed that Pol II binding frequency measured by SMF scales with enrichments determined using orthogonal measures such as ChIP-seq or PRO-seq (34). While this measurement is not time resolved, the frequency of each state in the cell population is a function of how often and how long each transcriptional intermediate occurs over time. Thus, quantification of the frequency of Pol II occupancy across species could enable us to identify the molecular steps that cause the difference in transcription kinetics between species.

Here, we used SMF to quantify PIC and Pol II occupancy at the promoters in mouse and *Drosophila* cell lines, two species with divergent transcription kinetic rates. While not observed in bulk assays, SMF data revealed that transcriptionally active mouse promoters are characterised by 3-5 times lower Pol II occupancy levels at the pausing site, than those in *Drosophila*. We applied mathematical modelling to infer Pol II kinetics at promoters from Pol II occupancy. To determine the relative contribution of each step of the process, we measured the changes in promoter-proximal Pol II levels upon inhibition of transcription initiation. We found that the turnover rate of Pol II at mouse promoters is higher than in *Drosophila*, but that these differences are insufficient to entirely explain the low occupancy levels observed in mouse cells. In turn, integration of these Pol II turnover rates into our model suggests that differences in the transcription initiation rate between species also significantly contribute to the differences in Pol II occupancy.

## Results

### Low amplitude Pol II footprints at mouse promoters

We have previously shown that SMF can be used to quantify the occupancy of nucleosomes, PIC, and Pol II at *Drosophila* promoters, and their dynamics across cell types (34). We now used a similar strategy to analyse a previously generated bait-capture SMF dataset in mouse embryonic stem cells (mESCs), that covers most of the annotated mouse promoters at high coverage (median of 100 molecules) (35). We first characterised the spatial distribution of the footprints created by the occupancy of the transcription machinery at core promoters of the mouse genome. We defined the TSS based on Cap Analysis of Gene Expression (CAGE) data (36). In case of multiple initiation sites are used (i.e. broad promoters in the mouse genome), we used the strongest CAGE peak (see Methods). We contrasted the average accessibility patterns at promoters of genes that are either highly active (top 5% Pol II ChIP-seq - Figure 1B-C) or inactive (bottom 10% Pol II ChIP-seq - Figure 1D). Chromatin accessibility was low at inactive promoters, while active promoters showed high accessibility upstream and a strongly phased +1 nucleosome down-stream of the TSS. Additionally, we subset TATA-containing promoters from the top 5% highly active promoters. The average profile of these TATA-containing promoters showed a strong footprint upstream of the TSS (Figure 1C), similar to what has been observed at *Drosophila* promoters (34, 37). We previously showed that this upstream footprint is created by PIC, as knockdown of TATA binding protein (TBP) attenuated this footprint in *Drosophila* (34). Thus, the occupancy profiles of mouse promoters generally recapitulate those of *Drosophila* promoters.

However, a direct overlay of the profiles from the two species revealed several differences (Figure 1E-F). First, the position of footprint corresponding to the +1 nucleosome downstream of the TSS is shifted by 20 bp in mouse promoters, in agreement with previous reports (38), and nucleosome positions determined by micrococcal nuclease digestion and sequencing (MNase-seq) (Figure 1E, middle panel). Second, while active *Drosophila* promoters harbour a prominent footprint downstream of the TSS at a position compatible with Pol II pausing (34), this footprint is absent when averaging signal over mouse promoters with comparable transcriptional activity (Figure 1E, upper panel). This difference in these downstream footprints is also visible, albeit less pronounced, when focusing on TATA-containing promoters where a low amplitude Pol II footprint is observed at mouse promoters (Figure 1F, upper panel). The observed difference in SMF footprints at the Pol II pausing site stands in contrast with the consistency in the accumulation of active RNA polymerase measured by bulk assays such as PRO-seq when performed in the same cell lines (Figure 1E-F, lower panel). We note however that the differences in SMF footprints are specific to Pol II, since the PIC footprint observable at TATA-containing promoters is almost identical between the two species (Figure 1F, upper panel). We next wondered if the observed differences may be a specific feature of mESCs that are in a pluripotent state. We compared the average profiles around the TSS of highly active genes in somatic cell lines representing various cell lineage (39). We observed very similar profiles in all the tested cell lines, with strong footprints for the PIC and low Pol II footprints downstream of the TSS (Figure S1A-B). This suggests that a low amplitude Pol II footprint is a general feature of mammalian promoters. Together, these observations suggest that the spatial patterns of promoter occupancy are globally conserved from *Drosophila* to mouse, but that Pol II occupancy levels may be reduced at mouse promoters.

### Quantification of the frequency of Pol II occupancy at mouse promoters

To quantify the frequency of promoter occupancy by the transcription machinery, we adapted the molecular classifier originally developed to study *Drosophila* promoters (34) to account for the shift in the Pol II and nucleosome positions relative to TSS in mouse (Figure S2A, see Methods for details). Based on the accessibility at four bins around the TSS, each DNA molecule is classified into different promoter states according to the presence or absence of a footprint at the positions occupied by the PIC and Pol II or the occupancy of the nucleosome at these positions (Figure S2B). With this strategy, we determined for each promoter the frequency of molecules in each of the promoter states (unassigned, nucleosome, unbound, PIC, PIC + Pol II, and Pol II). In total, we were able to quantify promoter state frequencies of 6,122 mouse promoters from the bait-capture SMF data. The resulting promoter state frequencies were consistent across replicates (Figure S2C), demonstrating that we sampled a sufficient number of DNA molecules covering each promoter. We further compared the frequency of the states with orthogonal measurements of Pol II or nucleosome occupancy at promoters. We found that the states were separated in two clusters corresponding to active and inactive promoter states (Figure S2D), consistent with our previous observations in *Drosophila* (34). Moreover, we observed a good agreement between the frequency of the states measured by SMF and the enrichment of the respective feature in bulk assays. This demonstrates the accuracy of our quantification of the occupancy by the transcription machinery and nucleosomes at single molecule resolution at mammalian promoters.

### Mouse promoters are characterised by low frequency of Pol II occupancy

Next, we compared the Pol II occupancies between species while accounting for their relative transcription activity. We ranked all promoters we were able to quantify with our method (mouse – 6,122 promoters; *Drosophila* – 5,912 promoters) by their activity, based on Pol II ChIP-seq signal at the promoters, and compared the distribution of promoter states between *Drosophila* and mouse (Figure 2A-D). At *Drosophila* promoters, an increase in promoter activity is correlated with a loss in nucleosome occupancy, gain in accessibility, and an increase in Pol II binding frequency (Figure 2A). Many highly-active promoters showed 20% of the molecules occupied by Pol II, as illustrated for the Svil promoter (17% - Figure 2B). At mouse promoters, we found similar loss in nucleosome occupancy and gain in accessibility as a function of Pol II ChIP-seq. Yet, while detectable, the Pol II state frequency was much reduced, with occupancies below 10% even at the most active TSSs (Figure 2C). For example, the Skp1 promoter is among the most active promoters but only 9% of the molecules is occupied by Pol II (Figure 2D).

**Fig. 2.**
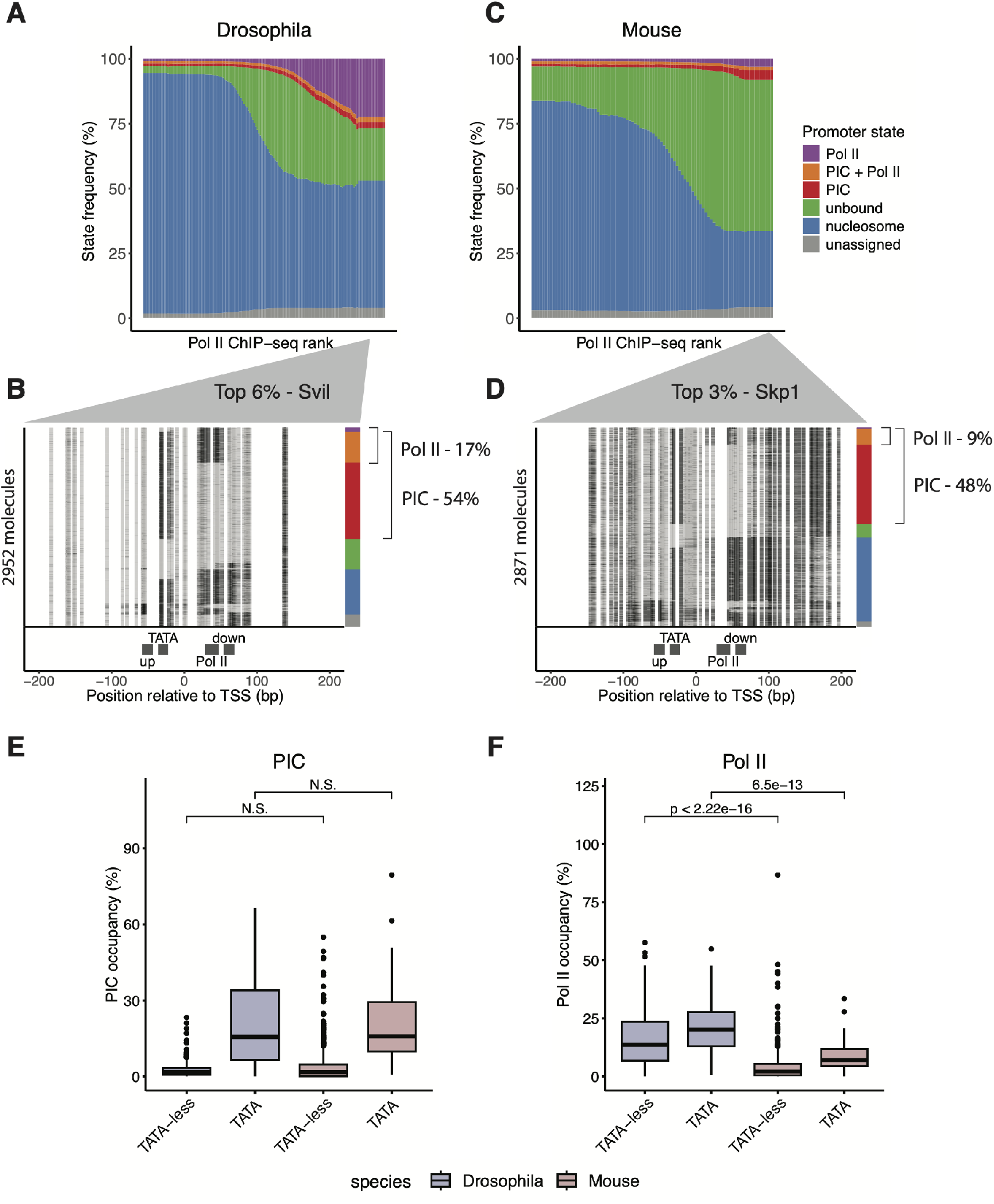
Lower Pol II binding frequency in mouse compared to *Drosophila* promoters. **(A)** Distribution of SMF-derived promoter state frequencies as a function of Pol II ChIP-seq level in *Drosophila* promoters (n = 5,912 promoters). Cumulative bar plot depicting the distribution of state frequencies. Promoters were binned based on log2 Pol II ChIP-seq signal, and the median frequency of each promoter state was calculated within each bin. Colour code represents each state as follows: purple – Pol II, orange – PIC + Pol II, red – PIC, green – unbound, blue – nucleosome, and grey – unassigned. **(B)** Single-locus example showing single-molecule sorting of a highly active, TATA-box containing *Drosophila* promoter (*Svil*, 94^th^ percentile by Pol II ChIP-seq). Each row in the single-molecule stack denotes a single DNA molecule and the methylation status of each cytosine in that molecule (methylated, accessible – light grey; unmethylated, protected – black). The positions of the four bins (upstream, TATA, Pol II, downstream) used for promoter state decomposition are shown below the single-molecule stack (see Methods). The vertical side bars on the right of the plot depict the frequency of each promoter state determined by single-molecule decomposition. The percentages of molecules harbouring footprints for the engaged Pol II and PIC are indicated on the right side of the plot. **(C** Distribution of SMF-derived state frequencies as a function of Pol II ChIP-seq level in mouse promoters (n = 6,122 promoters). Same representation as **(A). (D)** Single-locus example of a single-molecule sorting of a highly active, TATA-box containing mouse promoter (*Skp1*, 97^th^ percentile by Pol II ChIP-seq). Same representation as **(B). (E)** Comparison of PIC state frequency between mouse (TATA-less n = 572 promoters; TATA-containing n = 64 promoters) and *Drosophila* (TATA-less n = 422 promoters; TATA-containing n = 71 promoters) promoters. Boxplots represent the distribution of the frequency of PIC-bound molecules (PIC and PIC + Pol II states) at highly active promoters (top 5% Pol II ChIP-seq level). **(F)** Comparison of Pol II state frequency between mouse and *Drosophila* promoters. Boxplots represent the distribution of the frequency of Pol II-bound molecules (Pol II and PIC + Pol II states) at highly active promoters (top 5% Pol II ChIP-seq level). The analysis is stratified by the presence of a TATA-box at the promoter. The number of promoters included in this analysis is similar to that in **(E)**. The statistical comparisons between groups were performed using Wilcoxon rank-sum test. N.S. denotes non-significant.

To quantitatively estimate the magnitude of the difference in promoter occupancy between mouse and *Drosophila*, we looked at the distribution of PIC and Pol II occupancies across the top 5% highly active promoters that we were able to quantify using SMF (mouse – 636 promoters; *Drosophila* – 493 promoters). We stratified this analysis based on the presence or absence of the TATA-box, as in *Drosophila* TATA-containing promoters have a higher PIC and Pol II occupancy than TATA-less promoters (34). Consistent with *Drosophila*, the PIC occupancy is significantly increased at the TATA-containing promoters in mouse (Figure 2E). There are, however, no significant differences in the distribution of promoter occupancy by the PIC between the two species (Figure 2E). This is in contrast to the frequency of Pol II occupancy. The median Pol II binding frequencies at highly active *Drosophila* promoters are 14% (TATA-less) and 20% (TATA-containing), while those of mouse promoters are only 2% and 7%, respectively (Figure 2F). This suggests that Pol II occupancy at mammalian promoters is a low frequency event, occurring in <10% of the cells at any given time even at highly active promoters.

### Low frequency Pol II footprints are lost upon inhibition of transcription initiation

Low frequency events such as Pol II occupancy in mammals are harder to quantify by SMF, as it requires to sample more DNA molecules to observe them. To unambiguously confirm that the low frequency footprints captured downstream of the mouse TSS are created by Pol II, we measured the change in the Pol II occupancy, the frequency of molecules in PIC + Pol II and Pol II states, upon inhibition of transcription initiation. We used triptolide (TRP) at concentrations previously shown to deplete promoter-proximal Pol II in mESCs (40), and performed targeted SMF against a targeted set of 47 promoters selected to cover a diverse spectrum of Pol II occupancies and promoter structures (35 TATA-containing and 12 TATA-less promoters). This targeted approach generates thousands of single molecule measures for each locus, that are robust for precise and reproducible quantification of the low Pol II occupancy at mouse promoters. While Pol II occupancy in this set of promoters is generally low, ranging from 0% to 20% with a median of 2.7%, we obtained a consistent estimation across replicates (Figure S3A). We further confirmed that the promoter state frequencies obtained from the targeted SMF are in a high agreement with other orthogonal approaches (Figure S3B).

Upon inhibition of transcription using TRP, we observed a loss of the footprint downstream of the TSS of the *Amd1* gene, consistent with the loss of Pol II at the pausing site (Figure 3A, upper panel). In contrast, the footprint upstream of the TSS, created by the PIC, remained unaffected. Both of these observations were confirmed by quantifying the PIC and Pol II occupancy at single molecule level, showing no significant changes in the footprint PIC state (34% and 38%), while Pol II occupancy decreased from 12% to 1% (Figure 3A, lower panel). A reduction in the Pol II footprint was also evident at the TATA-less *Rsrp1* promoter, decreasing from 8% to 1% following TRP treatment (Figure S4). Extending this analysis to a larger set of 47 mouse promoters, we observed a consistent loss of Pol II occupancy upon blocking transcription initiation by TRP treatment in mouse promoters, similar to what has been observed in *Drosophila* promoters (34) (3B). Taken together, these results confirm that the low frequency footprints observed at mouse promoters are created by Pol II at the pausing site.

**Fig. 3.**
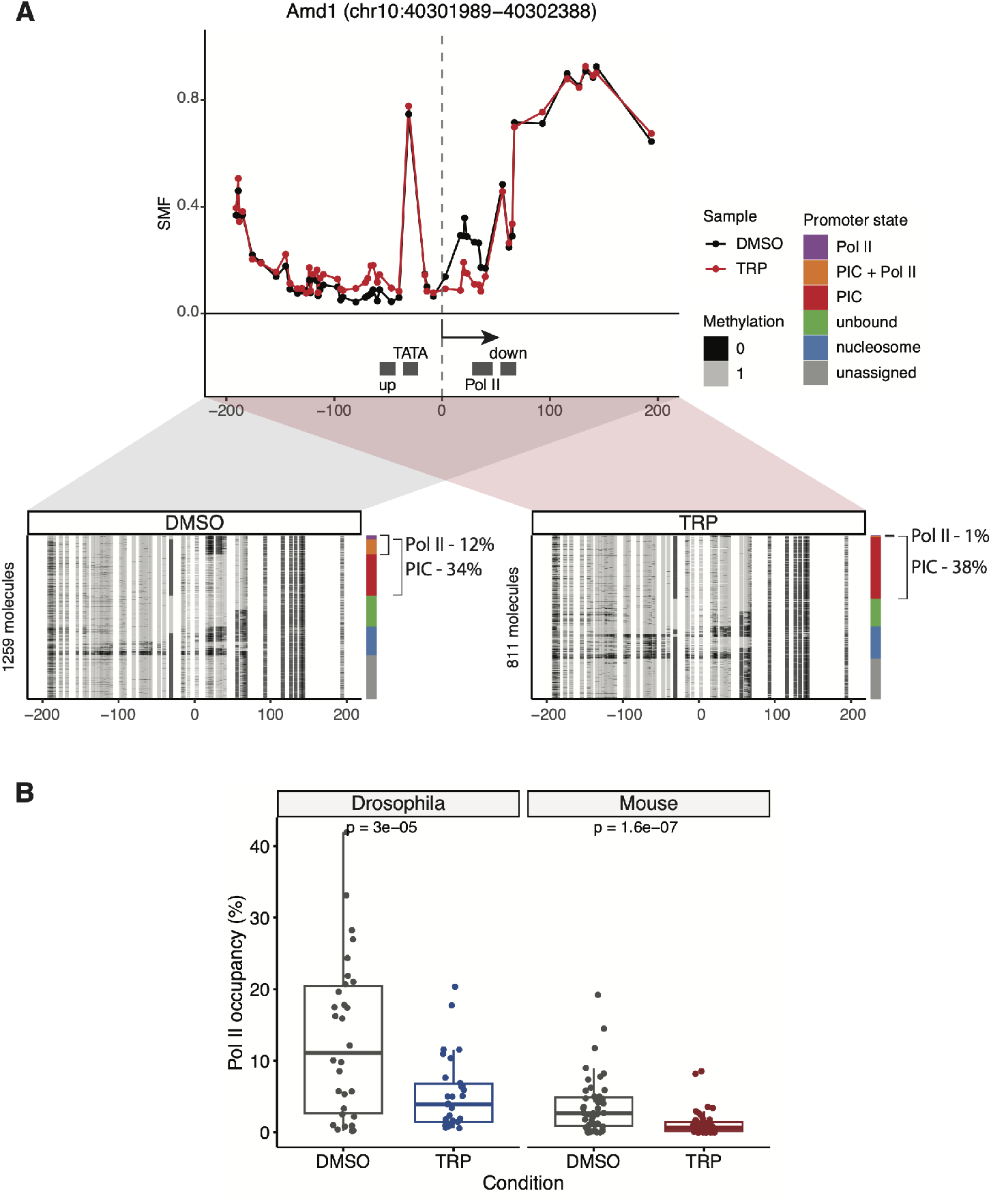
Inhibition of transcription initiation reduces Pol II footprint at mouse promoters. **(A)** Single-site example (*Amd1* promoter) shows a reduction in Pol II footprint upon triptolide (TRP) treatment. The experiment was conducted using targeted amplicon bisulfite sequencing on 47 selected promoters. The upper panel shows the average SMF plot (DMSO – black, TRP – red). The positions of the four bins (upstream, TATA, Pol II, downstream) used for promoter state decomposition are shown (see Methods). The x-axis represents the position relative to the TSS, while the y-axis shows the SMF signal (1 - methylation). The lower panel displays single-molecule stack plots for DMSO and TRP conditions. Each row denotes a single DNA molecule and the methylation status of each cytosine in that molecule (methylated, accessible – light grey; unmethylated, protected – black). The vertical side bars display the frequency of each promoter state. The percentages of molecules harbouring footprints for the engaged Pol II are indicated on the right side of the plot. **(B)** Loss of Pol II occupancy upon inhibition of transcription initiation in *Drosophila* (n = 30 promoters; (34)) and mouse promoters (n = 47 promoters). Boxplots represent the distribution of the frequency of Pol II-bound molecules (Pol II and PIC + Pol II states). Statistical comparisons between groups were performed using the Wilcoxon signed-rank test.

### A quantitative model to link molecular occupancy at core promoters with Pol II kinetics

We next aimed to identify the mechanisms explaining the interspecies differences in Pol II occupancy at promoters. Given the conservation of the general transcription machinery, we hypothesised that these differences might arise from variations in the kinetics of molecular processes, including the transcription initiation rate and Pol II turnover rate. Here, Pol II turnover rate is a combination of the rate at which Pol II undergoes premature termination and the rate at which Pol II progresses to elongation. To determine whether transcription initiation or Pol II turnover is more likely to explain the observed differences in Pol II occupancy, we developed a theoretical framework that relates promoter occupancy, either by nucleosomes or Pol II, to the effective rates of Pol II initiation and turnover. To do this, we implemented a stochastic model of transcription that explicitly considers three promoter states that can be captured by SMF: a closed state where the promoter is occupied by nucleosomes (*P*_*nuc*_); an unbound state consisting of accessible DNA without nucleosome or Pol II (*P*_*open*_); and a Pol II-bound state where the promoter is occupied by Pol II (*P*_*P olII*_) (Figure 4A). The transitions between these states are described by effective kinetic rates: nucleosome binding-unbinding (*k*_*c*_ and *k*_*o*_), transcription initiation (*k*_*i*_), and Pol II turnover (*k*_*t*_). Mechanistically the turnover rate (*k*_*t*_) is an effective rate that combines two distinct events that cannot be distinguished by our assay: Pol II entry into productive elongation or early transcription termination. This approach based on modelling SMF-derived states allows for a mechanistic interpretation of the kinetic parameters. This brings an improvement to existing stochastic models of transcription developed from imaging experiments where the kinetic parameters are not formally connected to precise molecular events (41, 42).

**Fig. 4.**
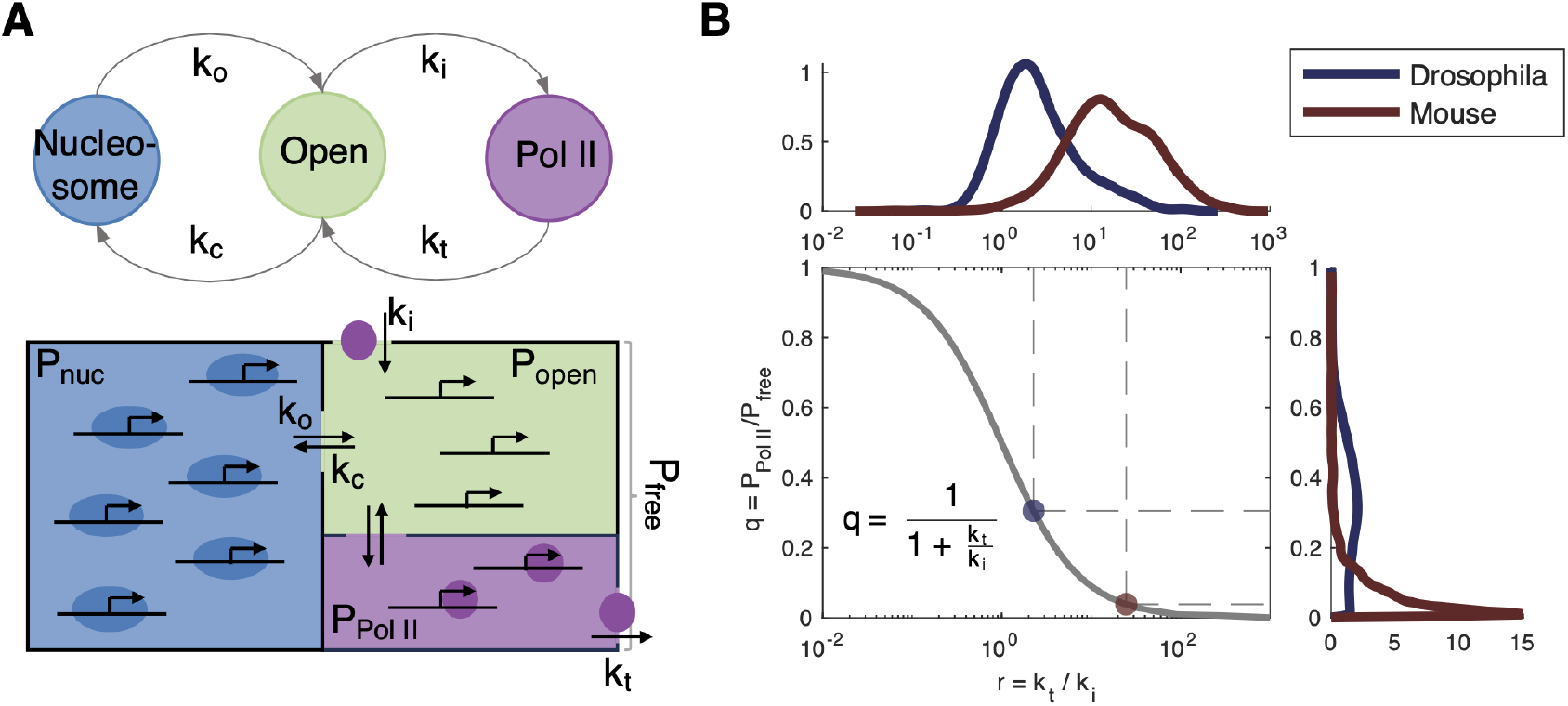
Kinetic modelling links Pol II occupancy with the initiation and Pol II turnover rates at promoters. **(A)** Schematic representation of promoter states derived from SMF experiments: “Nucleosome” indicates the promoter is bound by a nucleosome (blue), “Open” signifies that the promoter is unbound by either nucleosome or Pol II (green), and “Pol II” represents the promoter bound by Pol II (purple). Transitions between promoter states are described by four effective rates: nucleosome bound (*k*_*c*_), nucleosome unbound (*k*_*o*_), transcription initiation (*k*_*i*_), and Pol II turnover (*k*_*t*_). The probability of a promoter being in each state is a function of the rates (see Methods) and relates to the promoter frequencies observed by SMF. The probabilities of finding promoters in the Nucleosome (nucleosome state from SMF), Open (unbound and PIC states from SMF), and Pol II (PIC + Pol II and Pol II states from SMF) states are denoted as *P*_*nuc*_, *P*_*open*_, and *P*_*P olII*_, respectively. **(B)** Our kinetic model links the relative Pol II occupancy (*q*), defined as the ratio of DNA molecules bound by Pol II with respect to DNA molecules free of nucleosome (*q* = *P*_*P olII*_ */P*_*free*_), with the ratio of Pol II turnover rate (*k*_*t*_) to transcription initiation rate (*k*_*i*_) (see equation inside the figure). The relationship derived from the equation is depicted as a solid grey line. The y-axis displays the distributions of Pol II occupancies (*q*) as measured by genome-wide SMF experiments, highlighting a clear difference between *Drosophila* (blue) and mouse (red). The x-axis presents the predicted ratio (*r*) of the Pol II turnover rate (*k*_*t*_) and the initiation rate (*k*_*i*_) for each promoter. The distributions of these predicted ratios are presented on the top, illustrating a clear difference between *Drosophila* (blue) and mouse (red). The dots represent median values of (*q*) and (*r*) for *Drosophila* (blue) and mouse (red).

Solving our model at steady state leads to simple expressions for the expected frequencies of each promoter state as a function of the kinetic rates (see Methods). In this simple model, the relative Pol II occupancy (*q*), defined as the ratio of DNA molecules bound by Pol II with respect to DNA molecules free of nucleosome, depends only on the ratio of the Pol II turnover rate (*k*_*t*_) and the initiation rate (*k*_*i*_) through a simple sigmoid function (Figure 4B). In other words, the Pol II occupancy is the result of the equilibrium between how fast the Pol II is loaded at the promoter and how fast it leaves (to either elongation or early termination). By integrating the promoter state frequencies estimated from SMF data into our model, we can estimate the ratio between rates (*k*_*t*_*/k*_*i*_) for each promoter in the *Drosophila* and mouse genome (Figure 4B). We observed a 7-fold difference in the median of this ratio between species (Figure 4B). In summary, we build a theoretical model that allows us to relate changes in Pol II frequency with the kinetic rates of the different steps of the process.

### Faster Pol II turnover and lower initiation at mouse promoters

Our model predicts that the differences in Pol II occupancy between species could arise from a higher transcription initiation rate in *Drosophila* or a higher Pol II turnover rate in mouse; or a combination of both. Our model quantitatively links the occupancy as measured by SMF to the ratio of the initiation rate (*k*_*i*_) and the Pol II turnover rate (*k*_*t*_). Thus, there is an interdependence between these parameters, allowing us to predict one if the other one can be measured. For instance, we would be able to estimate the initiation rate if we can measure the turnover rate of Pol II at promoters.

To estimate Pol II turnover rates in each specie, we treated *Drosophila* S2 cells and mESCs with TRP to inhibit transcription initiation and measured the change in promoter-proximal Pol II at multiple time points (0, 2.5, 5, 10, and 20 min) using PRO-seq (43) (see Methods). The PRO-seq intensity and their changes over time were highly correlated between the two replicates (Figure S5). We found that the turnover of promoter Pol II following the inhibition of transcription initiation is faster in mouse than in *Drosophila* (Figure 5A). For example, although the mouse *Polr2a* promoter and *Drosophila Fur1* promoter have comparable Pol II binding frequencies measured by SMF (19%), the *Polr2a* promoter showed a faster decay of promoter-proximal Pol II than the *Fur1* promoter (Figure 5B-C). To further investigate the differences in Pol II turnover between species, we clustered promoters according to the changes in promoter-proximal Pol II measured by PRO-seq over time (Figure 5D-E). We fitted an exponential decay model on each cluster to estimate the half-life of Pol II at promoters. We observed that about 20% of *Drosophila* promoters have a half-life of less than 5 minutes (Figure 5D). In contrast, more than 80% of mouse promoters have a half-life of less than 5 minutes, suggesting that Pol II turnover rate is generally faster at mouse promoters (Figure 5E).

**Fig. 5.**
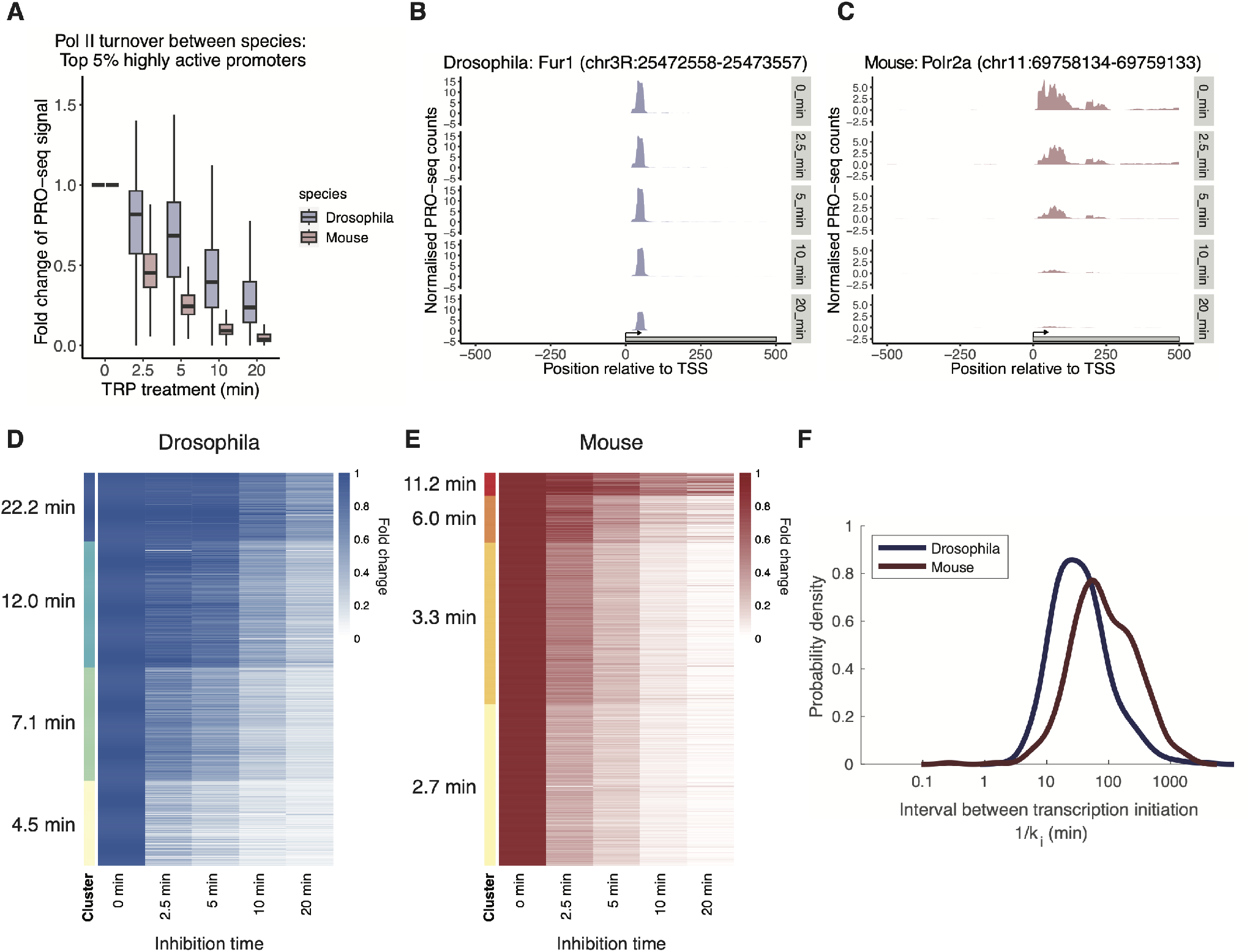
Mouse promoters exhibit a faster Pol II turnover rate and a lower initiation rate compared to *Drosophila* promoters. **(A)** Comparison of Pol II turnover following inhibition of transcription initiation via time-course triptolide (TRP) treatment between the top 5% highly active *Drosophila* (n = 930 promoters) and mouse promoters (n = 1,254 promoters). PRO-seq reads were counted around the TSS [-100:200]. PRO-seq reads were then normalised for each sample using Spike-in read counts (see Methods). Each data point is an average of the two replicates, except in the 2.5 min TRP treatment, where only one replicate passed quality control. For each promoter, the fold change of normalised PRO-seq reads was calculated at each time point relative to the DMSO treatment (0 min). The x-axis indicates the duration of TRP treatment, while the y-axis represents the fold change of normalised PRO-seq counts compared to the DMSO treatment. **(B-C)** Single-site example of time-course PRO-seq data following the time-course TRP treatment from **(B)** the mouse *Polr2a* promoter and **(C)** the *Drosophila Fur1* promoter, both with a similar initial Pol II binding frequency (19%). Normalised PRO-seq counts were smoothed over 25 bp. **(D-E)** K-means clustering of the fold changes in normalised PRO-seq counts at each time point relative to the DMSO treatment (0 min) for **(D)** the top 5% *Drosophila* promoters and **(E)** the top 5% mouse promoters. The half-life of Pol II for each cluster was determined by fitting an exponential decay model to the normalised PRO-seq counts and shown next to the heatmap. **(F)** Modelling of relative Pol II occupancy (*q*) and Pol II turnover rate (*k*_*t*_) reveals the difference in transcription initiation rate (*k*_*i*_) between mouse and *Drosophila*. The density plot is shown with x-axis represents interval between transcription initiation, which is the inverse of the initiation rate (1*/k*_*i*_).

We then integrated these experimentally determined Pol II turnover rates into our model to ask if these are sufficient to explain the differences in Pol II occupancy observed between species. Using the interdependency between the initiation and the turnover rates, we predicted the initiation rates (*k*_*i*_) for each gene as a function of Pol II occupancy (*q*) and Pol II turnover (*k*_*t*_) (see Methods). If the turnover rates were sufficient to explain the differences in occupancy, the distribution of initiation rates should be similar between species. In contradiction with this hypothesis, we observed that the predicted initiation rates are significantly faster for *Drosophila* promoters (median = 0.032 min^-1^) than those of mouse promoters (median = 0.014 min^-1^) (Figure 5F). This means that a Pol II molecule initiates on average every 32 minutes at *Drosophila* promoters, while it initiates every 72 minutes at mouse promoters. These slower initiation rates in mouse are consistent with the general notion of lower burst frequency in mammalian cell lines compared to *Drosophila* from live transcription imaging experiments (15, 16). Together, our results suggest that while Pol II turnover contributes to the differences in Pol II occupancy between species, it is not sufficient to entirely explain them. The remaining fraction of the differences could be explained by different rates of transcription initiation.

## Discussion

Here, we found that promoter-proximal Pol II pausing at mouse promoters is much less frequent than at *Drosophila* promoters. While up to 40% of the cells have Pol II engaged at active genes in *Drosophila*, this drops to <10% at most promoters of active genes in mouse cells. This difference in Pol II occupancy contrasts with the occupancy of other factors, such as the PIC, which shows comparable levels of occupancy at promoters in both species. Mechanistically, we show that these contrasts in occupancy reflect differences in the kinetics of recruitment and turnover of Pol II at the promoter. Mouse cells are characterised by faster turnover of Pol II and a lower initiation rate, which together reduce the number of Pol II molecules engaged on DNA at any given time (Figure 6).

**Fig. 6.**
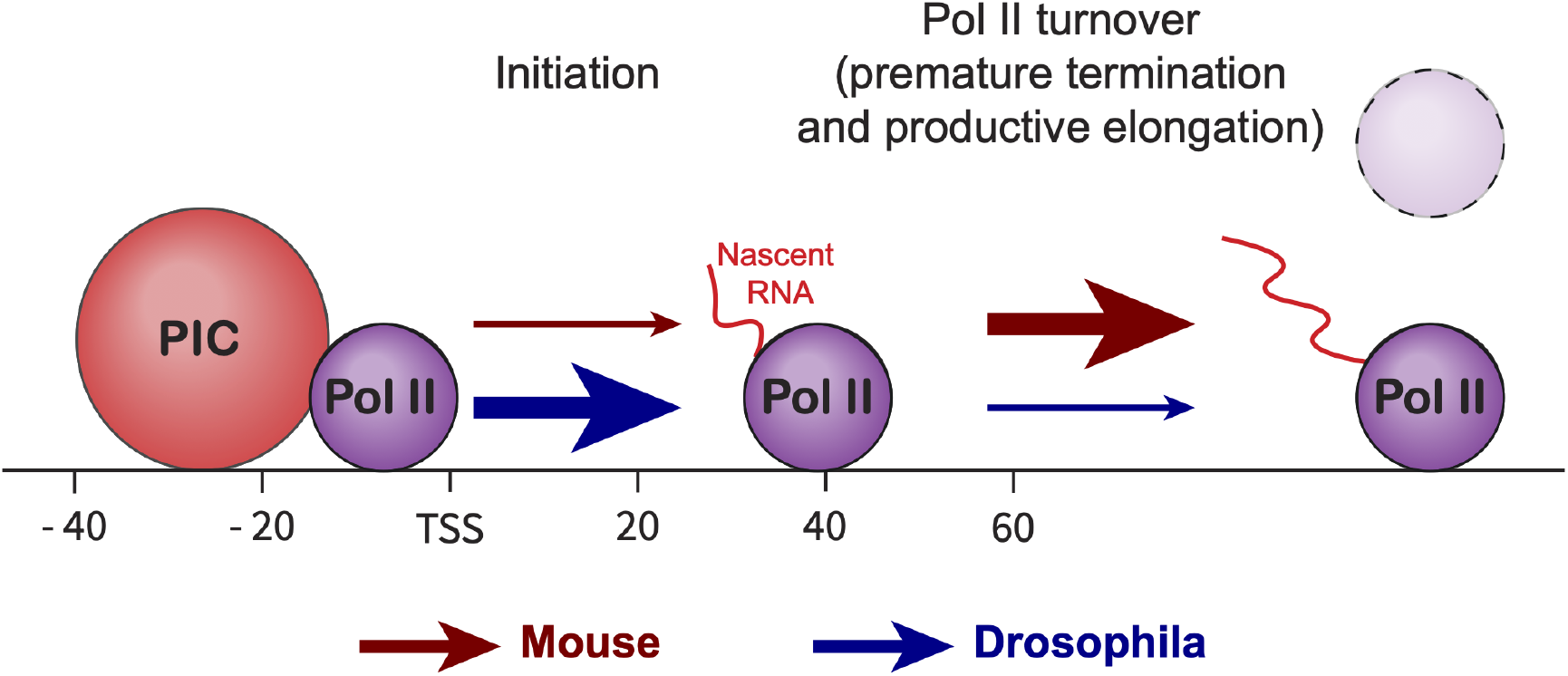
Summary of the differences in Pol II occupancy kinetics between mouse and Drosophila promoters. The lower Pol II occupancy at mouse promoters can be attributed to 1) a reduced transcription initiation rate and 2) an increased Pol II turnover, which includes both premature termination and productive elongation, in comparison to *Drosophila* promoters. The colours of the arrows represent mouse (red) and *Drosophila* (blue), with the size of the arrows indicating the rate of activity – smaller arrows represent lower rates, while larger arrows denote higher rates.

*Drosophila* and mouse cells are popular experimental models for studying transcription mechanisms. Pol II occupancy has been profiled with various genomics assays such as ChIP-seq, PRO-seq or sequencing of short capped transcripts (32, 44, 45). It is thus surprising that these differences in Pol II occupancy were not detected earlier. This may arise from fundamental limitations of bulk assays to estimate global changes. Bulk assays are based on the experimental enrichment of the Pol II-associated molecules. In contrast, SMF measures all the DNA molecules of a cell population, regardless of their occupancy by Pol II. This fundamental difference in experimental principles provides a unique opportunity to quantify the frequency of factor occupancy on DNA, which cannot be estimated from bulk genomics data (34, 46). This approach can, in principle, be expanded to any DNA-associated factors for which reference occupancy data exist (33).

Regulation of transcription after initiation through Pol II pausing has been proposed to be critical for the regulation of genes requiring rapid induction, such as those involved in key developmental transitions or stress responses (32, 47, 48). The observation of the characteristic accumulation of Pol II downstream of the TSS led to the model that the rapid transcription response would be mediated through the release of these preloaded polymerases into the gene body (49). Several lines of evidence from genomics and live-cell-imaging approaches argue that Pol II at the pause site rapidly turns over at a large fraction of genes (34, 50–54), suggesting an alternative model where paused genes frequently experience early transcription termination. The identification and characterisation of early transcription termination by the Integrator complex provided a mechanism supporting this model (55–58). Our data suggest that in mammals, Pol II is detected at the paused site in a very small fraction of cells. Moreover, we found that Pol II turnover was faster at mouse than *Drosophila* promoters, with half-lives of less than 5 minutes in most highly active mouse promoters. Our findings are in agreement with previous studies conducted in human cells (51, 52), suggesting that rapid turnover of promoter-proximal Pol II is a characteristic of mammalian promoters. In these conditions the pool of polymerases available for rapid release in the gene body is unlikely to be sufficient to explain the rapid and sustained induction of paused genes. Together these data further support of a model where Pol II pausing primarily occurs through rapid cycles of initiation and early termination. In this model, pause release occurs through a change in the fate of polymerase from active termination to productive elongation. The precise mechanisms underlying this switch and its specificity remain to be identified.

By integrating experimental data into our quantitative model, we found that the Pol II turnover rate alone cannot fully account for the discrepancy in Pol II occupancy between mouse and *Drosophila*. We predict that the remaining variance is due to a lower transcription initiation rate at mouse promoters. This prediction is in agreement with the observation that transcriptional bursts at mammalian genes are interspersed with long periods of transcriptionally inactive states (12–16). The presence of TATA box has been associated with longer durations of the bursts and more frequent Pol II initiation (4, 10). Our data are in line with this notion, as Pol II occupancy is higher at TATA-containing promoters compared to TATA-less promoters in both mouse and *Drosophila*. Transcriptional bursting has been described by models such as the two-state telegraph model (3, 41, 59), as well as more intricate models that accommodate complex molecular mechanisms underlying transcription, including Pol II pausing (10, 60). A limitation of these models is that they provide only phenomenological descriptions of the process, but the underlying molecular mechanisms are not explicitly identified. Here we show that single-molecule assays such as SMF used in this study, can be used to formally link occupancies of factors like nucleosomes, PIC, and Pol II at *cis*-regulatory elements with kinetics of regulatory processes. Similar principles have recently been applied to relate TF binding with promoter state and transcription at synthetic cis-regulatory elements (61), demonstrating the potential of the approach to connect the molecular states of regulatory elements with transcriptional kinetics.

## Methods

### Cell culture and treatment

DNA methyltransferase triple knockout mouse embryonic stem cells (TKO mESCs) (62) were cultured at 37°C on 10 cm plates coated with 0.2% gelatin. The cells were maintained in Dulbecco’s Modified Eagle Medium (DMEM, Thermo Fisher) supplemented with 15% fetal bovine serum (FBS, Sigma-Aldrich), 1% sodium pyruvate (Sigma-Aldrich), 1 mM L-glutamine, 1× non-essential amino acids (NEAA, Thermo Fisher), 0.2% leukemia inhibitory factor (LIF, produced in-house), and 0.01% β-mercaptoethanol. Media were regularly replaced, and cells were passaged as needed to maintain optimal growth conditions.

Schneider-S2 cells were grown at 25°C in Schneider’s *Drosophila* medium (LifeTech: 21720-001) supplemented with 10% FBS.

To inhibit transcription initiation, TKO mESCs were treated with 500 nM triptolide (TRP) (Sigma) for 30 minutes. Control cells were treated with an equal volume of DMSO. Following treatment, both control and TRP-treated cells were prepared for amplicon Single Molecule Footprinting (SMF). For the time-course TRP experiment, TKO mESCs and S2 cells were treated with 10 µM TRP for varying durations (2.5, 5, 10, and 20 minutes). Control cells were treated with an equivalent volume of DMSO for 20 minutes. After treatment, cells were harvested for PRO-seq.

### Single Molecule Footprinting

Dual-enzyme SMF was performed following the protocol from (33, 35). Briefly, cells were collected by trypsinization, washed in ice-cold PBS and counted with a trypan blue based automatic counter (Bio-Rad). Then, per SMF reaction, 250,000 cells were lysed in chilled lysis buffer (10 mM Tris (pH = 7.4), 10 mM NaCl, 3 mM Mgcl2, 0.1 mM EDTA, 0.5% NP40) for 10 minutes on ice. Nuclei were pelleted by centrifuging 5 minutes at 4°C and 1000 x g, and washed in wash buffer (10 mM Tris (pH = 7.4), 10 mM NaCl, 3 mM MgCl2, 0.1 mM EDTA). Next, nuclei were resuspended in 1x M.CviPI reaction buffer (50 mM Tris (pH 8.5), 50 mM NaCl, 10 mM DTT and 300 mM sucrose) and incubated for 7.5 minutes at 37°C with 200U of M.CviPI (NEB M0227L) and 0.6 mM SAM (NEB). A second 7.5 minute incubation was performed with an additional 100U of M.CviPI and 0.128 mM SAM. Next, 10 mM MgCl2, 0.128 mM SAM and 60U of M.SssI (NEB-M0226L) were added, and nuclei were incubated a third round of 7.5 minutes at 37°C. The methylation reaction was stopped with stop buffer (20 mM Tris, 600 mM NaCl, 1% SDS 10 mM EDTA). All samples were digested with Proteinase K (200 mg/ml) overnight at 55°C, followed by phenol/chloroform purification and isopropanol precipitation of DNA.

### Amplicon SMF

A set of 62 mouse promoters of protein coding genes were targeted with amplicon-SMF. These promoters were selected to cover promoters with and without a TATA box, covering a variety of expression levels (by RNA-seq) and activity levels (Pol II ChIP at the promoter). Bisulfite specific primers were designed using a custom R script based on using Primer3 with slight modifications, designing against in silico bisulfite converted sequences excluding CpG and GpC dinucleotides. For some promoters, two primer-pairs were designed for control purposes. Additionally, the primer plate covered 8 tRNA promoters (not analysed here) and re-used 12 primer-pairs used before for control purposes: 6 covering promoters, 6 covering TF binding sites (35).

Amplicon SMF was performed as previously described (33, 35). Briefly, the isolated footprinted DNA (1 µg per sample) was bisulfite converted with the Epitect bisulfite conversion kit (QIAGEN), according to manufacturer’s instructions. The converted DNA was distributed over a 96 wells plate, and PCR amplified in a total volume of 16 µL with 1x KAPA HiFi HotStart Uracil+ ReadyMix (Roche) and 625 nM primers (one primer-pair, forward and reverse combined), using the following cycling protocol: 3 minutes at 95°C, 35 cycles of (20s at 98°C, 30s at 56°C, 60s at 72°C), 5 minutes at 72°C. PCR product was checked by gel electrophoresis, 10 µL per reaction was pooled and purified using 0.8x AMPureXP bead purification (Beckman Coulter). 1 µg of purified DNA per sample was used to prepare sequencing libraries, using the NEBNext Ultra II Kit according to manufacturer’s protocol and using NEBNext Multiplex Oligos to multiplex up to 12 libraries. Libraries were sequences on a MiSeq instrument to generate 250bp paired end reads.

### PRO-seq

Precision run-on sequencing (PRO-seq) was performed as previously described (43, 44) with slight modifications. For the *Drosophila* S2 cell samples, 10 million *Drosophila* S2 cells were used per sample with a spike-in of 0.1 million (1%) TKO mESCs. For the TKO mESC samples, 5 million TKO mESCs were used per sample with a spike-in of 0.05 million (1%) *Drosophila* S2 cells. Two replicates per TRP treatment time point were generated, with each replicate prepared simultaneously for all cell lines and conditions. Cells were harvested using Trypsin-EDTA on ice and washed twice with 10 mL ice-cold PBS, followed by centrifugation at 1000x g for 4 minutes at 4°C. Permeabilisation was performed in 10 mL of permeabilisation buffer (10 mM Tris-HCl pH 8.0, 250 mM sucrose, 10 mM KCl, 5 mM MgCl2, 1 mM EGTA, 0.1% Igepal, 0.05% Tween-20, 0.5 mM DTT, 10% (vol/vol) glycerol, 1 tablet of protease inhibitor per 50 mL, 4 units/ml SUPERaseIN inhibitor) on ice for 10 minutes. The cell pellets were then washed twice with 10 mL of ice-cold cell wash buffer (10 mM Tris-HCl pH 8.0, 250 mM sucrose, 10 mM KCl, 5 mM MgCl2, 1 mM EGTA, 0.5 mM DTT, 10% (vol/vol) glycerol, 1 tablet of protease inhibitor per 50 mL, 4 units/mL SUPERaseIN inhibitor) with centrifugation at 1000x g for 4 minutes at 4°C. Cells were resuspended in 50 µL freeze buffer (50 mM Tris-HCl, pH 8.0, 40% (vol/vol) glycerol, 5 mM MgCl2, 1.1 mM EDTA, 0.5 mM DTT and 4 units/mL SUPERaseIN inhibitor), snap-frozen, and stored at -80°C until further use. The nuclear run-on reaction was performed as a 2-biotin run-on. For this, 50 µL of the 2x Nuclear-run on master mix (10 mM Tris-Cl pH 8.0, 5 mM MgCl2, 1 mM DTT, 300 mM KCl, 40 µM Biotin-11-UTP, 40 µM Biotin-11-CTP, 40 µM ATP, 40 µM GTP, 1% Sarkosyl, 1 µL SUPERaseIN inhibitor) was prepared for each sample. The reaction mix was pre-heated at 37°C (30°C for *Drosophila* S2 cell samples) and 50 µL of the pre-calculated number of cells were added to each reaction vial, mixed thoroughly and incubated at 37°C (30°C for *Drosophila* S2 cell samples) for 5 minutes with shaking (750 rpm). To stop the reaction, 350 µL of the RL Buffer from the Norgen RNA extraction kit were added and vortexed. 240 µL of 100% ethanol was added to the mixture and vortexed again. RNA extraction was performed according to the kit’s manual. The final RNA was eluted twice with 50 µL H2O and pooled to a final volume of 100 µL. For the base hydrolysis, the RNA was denatured for 30 seconds at 65°C and snap-cooled on ice. 25 µL of ice-cold 1N NaOH were added and incubated on ice for 10 min. RNA was precipitated by adding 125 µL Tris-HCl (pH 6.8), 5 µL NaCl, 1 µL GlycoBlue and 650 µL 100% EtOH and centrifugation at 20,000x g at 4°C for 20 minutes. The RNA pellet was washed with 70% EtOH, air-dried and resuspended in 6 µL H2O. For the 3’ RNA adaptor ligation, 1 µL of REV3 3’RNA adaptor dilution (10 µM), heat-denatured at 65°C for 20 seconds and placed on ice, was added to the 6 µL resuspended RNA. 13 µL of the RNA ligation mix using T4 RNA ligase I was added and incubated at 25°C for 1 hour. Biotin RNA enrichment was performed by adding 55 µL binding buffer (10 mM Tris-HCL pH 7.4, 300 mM NaCl, 0.1% Tergitol, 1 mM EDTA) to each ligation reaction, followed by 25 µL of pre-washed streptavidin beads. Reactions were incubated on a rotator set to 8 rpm at room temperature for 20 minutes. Beads were washed once with ice-cold high-salt buffer (50 mM Tris-HCL pH 7.4, 2M NaCl, 0.5% Tergitol, 1mM EDTA), transferred to new tubes, and washed once with low-salt buffer (5 mM Tris-HCL pH 7.4, 0.1% Tergitol, 1mM EDTA). On-bead 5’ hydroxyl repair was performed by resuspending the beads in 19 µL of the PNK mix and incubated at 37°C for 30 minutes with soft shaking (350 rpm). For the 5’ cap repair reaction, the beads were resuspended in the enzyme mix containing RppH and ThermoPol Reaction Buffer and incubated at 37°C for 1 hour with soft shaking (350 rpm). On-bead 5’ RNA adaptor ligation was performed by resuspending the beads in 7 µL of REV5 5’RNA adaptor dilution (6 µL of H2O + 1 µL of 10 µM REV5), heat-denatured at 65°C for 30 seconds and placed on ice. 12 µL of the RNA ligation mix using T4 RNA ligase was added and incubated at 25°C for 1 hour. Beads were washed once with ice-cold high-salt buffer, transferred to new tubes, and washed once with low-salt buffer. The RNA was cleaned-up using Trizol and chloroform. The RNA was reverse transcribed using the Maxima H minus RT enzyme and the RP1 reverse-transcription primer. The correct number of PCR cycles was determined by test PCR and Bioanalyzer analysis. In the end, the final PCR was performed using 13 cycles and the final library was cleaned-up using magnetic SPRI beads at a ratio of 1.8x and further size selected with a SPRI beads ratio of 1x to remove primer dimers. The library was run on a NextSeq 2000 P3 with 50 bp single-end sequencing.

### Sequencing data preprocessing

A list of sequencing data used in this study is shown in the Table S1. SMF data was processed as previously described (33, 35). Briefly, raw paired-end sequencing reads were pre-processed using Trim Galore! (version 0.6.7) (63) to remove unpaired and low-quality reads, trim Illumina adapter sequences, and trim lowquality bases. The whole-genome *Drosophila* SMF data were pre-processed similarly using Trim Galore!, trimming one extra base from the start of each read to allow proper alignment of short fragments. The trimmed reads were aligned against a bisulfite index of the *Mus musculus* (mm10) or *Drosophila melanogaster* (dm6) genome using the R package QuasR (64), which uses Bowtie (65) as an aligner, with specific alignment parameters (alignmentParameter = -e 70 - X 1000 -k 2 –best –strata) and keeping only uniquely aligned reads. Duplicated reads were removed for genome-wide and bait-capture, but not for amplicon SMF data, using the tool MarkDuplicates from Picard (version 2.15.0) (66).

For PRO-seq data generated in this study, raw sequencing reads were trimmed using Cutadapt (version 3.5) (67). The UMIs were detected and extracted using UMI-tools (version 1.1.2) (68). Reads were aligned to a normal index of the respective genome of the sample and spike-in (mm10 and dm6 for TKO mESC samples and *Drosophila* S2 cell samples, respectively) using QuasR with default alignment parameters. Aligned reads were deduplicated using UMI-tools. Reads aligned to rRNAs were removed.

Other publicly available datasets (ChIP-seq, RNA-seq, MNase-seq, PRO-seq) were trimmed using Trim-Galore and aligned using QuasR against a normal index of the respective genome (mm10 or dm6) with default alignment parameters.

### SMF - Single molecule methylation call

SMF data analysis was performed using the R package SingleMoleculeFootprinting (‘promoter’ branch) (33). Methylation was called on all cytosines in aligned reads with a minimum bisulfite conversion rate of 80%, using the CallContextMethylation function, which was built on the QuasR function qMeth (64). For single-enzyme SMF (Figure S1), only Cs in the DGCHN (D = no C, H = no G, N = any base) were considered. For dual-enzyme SMF (Figures 1-3), all methylation information from CpG and GpC contexts were used.

### SMF - Single molecule sorting

For single molecule analysis, the molecular classifier from our previous study (34) was adapted to adequately assess the presence of promoter-proximal footprints at mouse promoters. In brief, four bins of interest were defined relative to the TSS. The bins include: upstream [-58:-43], TATA-box [-36:-22], Pol II [28:47], and downstream [54:69] (Figure S2A). For each TSS, all reads covering at least one relevant C per bin in all four bins were analysed. Thus, the analysis excluded any TSSs where any bin did not contain at least one relevant C. For each read, the average methylation per bin was calculated and rounded to 0 or 1, creating a 4 bits vector classifying the state of every read among 24 = 16 theoretical possibilities (Figure S2B). The reads were grouped by the methylation pattern within four bins into six promoter states: Pol II, PIC + Pol II, PIC, unbound, nucleosome, and unassigned (Figure S2B). For whole-genome *Drosophila* S2 cells and bait-capture TKO mESCs, the frequencies of promoter states were calculated per replicate for each TSS with at least 20 sorted reads. The promoter state frequencies were then averaged across the replicates. For amplicon SMF, only TSSs with at least 100 sorted reads (47 promoters) were retained for the analysis. The frequencies of promoter states were calculated per replicate and then averaged across the replicates. For the analysis in Figure 2E-F and Figure 3B, the frequencies of Pol II and PIC + Pol II were combined as a Pol II binding frequency. Similarly, the frequencies of PIC and PIC + Pol II were combined as a PIC binding frequency.

### SMF - Plots

For composite plots (Figure 1B-D, Figure S1), the average SMF (1 - % methylation) is displayed per C as a dot, along with the average over a 20 bp window as a line. In Figure 1E-F, only the average line is displayed.

Functions from the R package SingleMoleculeFootprinting (‘promoter’ branch) (33) were used for generating single locus plots (Figure 2B, Figure 2D, Figure 3A, Figure S4).

### Annotation of TSSs and promoters

TSSs were defined by taking the starting positions of all autosomal RefSeq transcripts (mm10 and dm6). Next, these TSSs were refined by shifting them to the strongest CAGE-seq peak within 50 bp of the TSS on the same strand as the gene (for mouse: CAGE peaks from the FANTOM5 project (69), lifted over to mm10 and reannotated by (36); for *Drosophila*: CAGE peaks from the modENCODE project (70), lifted over to dm6 by us). For TSSs without a corresponding CAGE-seq peak, the original TSSs were retained. After this step, duplicated start sites were removed. The presence of a TATA motif was determined by searching for the consensus motif (TATAWAWR) in its theoretical position relative to the TSS ±5 bp [-37:-18]. Promoter activity (e.g. top 5% highly active) was ranked according to Pol II ChIP-seq data (see below).

### Comparison with external datasets

For Pol II ChIP-seq, the start of each read was shifted 75 bp downstream to account for the average fragment size. The amount of Pol II was determined by counting the reads in a window of [-200:100] around each TSS, using the QuasR function qCount (64).

For RNA-seq, RPKM values were calculated per Ref-Seq transcript and assigned to the corresponding TSS. Because PRO-seq is strand-specific and the nascent RNA is sequenced from the 3’ end, only reads aligning to the strand opposite the gene were assessed. The amount of Pol II was determined by counting the reads in a window of [-200:100] around each TSS.

For the mouse MNase-seq data, the start of each read was shifted 80 bp downstream to account for the average fragment size. For the paired-end *Drosophila* MNase-seq data, the centre of each mapped fragment was used. The number of mapped reads was counted in a window of [-40:30] around the TSS to include only the nucleosomes overlapping the promoter and to exclude the -1 and +1 nucleosomes. The positions of nucleosomes and Pol II relative to the TSS (Figure 1E-F) were determined using the QuasR function qProfile on MNase-seq and PRO-seq data, respectively. In the window of [-250:250] around each TSS, read counts were normalised to reads per million mapped reads (RPM) and smoothed over a 20 bp window. RPM values at each position around the TSS were averaged across the top 5% highly active promoters (Figure 1E) or the top 5% highly active, TATA-containing promoters (Figure 1F). To enable a comparison of MNase-seq and PRO-seq data from two different species in the same plots, the average RPM for PRO-seq and MNase-seq at each position around TSS was normalised to the Z-score.

The global relationship between SMF-derived promoter states, Pol II ChIP-seq, RNA-seq, PRO-seq, and MNase-seq were assessed using Spearman correlation (Figure S2D, Figure S3B).

### PRO-seq analysis

PRO-seq data in this study generated from the time-course TRP treatment experiment (see above for details). Two samples were dropped (one replicate of TKO mESCs at 2.5 minutes and one replicate of *Drosophila* S2 cells at 2.5 minutes) and as they did not pass the quality control.

After alignment against the respective genomes (mm10 or dm6), PRO-seq reads were counted in a window around TSSs [-100:200] using the QuasR function qCount (64). Because PRO-seq is strand-specific and the nascent RNA is sequenced from the 3’ end, only reads aligning to the strand opposite the gene were assessed. For the correlation analysis between replicates (Figure S5), PRO-seq counts at the TSSs were normalised to RPM and compared. Under the assumption that read counts from spike-ins should be constant across samples, spike-in reads were used to perform inter-sample normalisation and enable quantification of the global effects expected to occur upon inhibition of transcription (Figure 5). For TKO mESC samples, PRO-seq read counts from each sample were normalised by the total reads mapped against the dm6 genome within the same sample. Similarly, for *Drosophila* S2 samples, PRO-seq read counts from each sample were normalised by the total reads mapped against the mm10 genome within the same sample. Replicates were averaged, and fold changes compared to the DMSO control were calculated (Figure 5). The analysis was restricted to highly active genes (top 5% Pol II ChIP-seq as defined above). To determine the Pol II turnover rate (Figure 5D-E), fold changes of PRO-seq signal at top 5% highly active promoters were clustered using k-means clustering (k = 4). For each cluster, an exponential decay model was fitted to the normalised PRO-seq counts, assuming a Poissonian noise model. Turnover rates (*k*_*t*_) were estimated by maximizing the likelihood, and confidence intervals were approximated by calculating the Hessian at the maximum. Half-lives where then calculated using the following expression *t*_1*/*2_ = *log*(2)*/k*_*t*_. This analysis revealed the half-life of promoter-proximal Pol II for each gene cluster.

### Mathematical model linking promoter occupancy state with transcriptional bursting

The simplest stochastic model widely used to describe transcriptional bursting is the so-called telegraph model (13, 41, 59), in which the gene can exist in two states, ‘ON’ and ‘OFF,’ with transcription occurring at a certain initiation rate in the ‘ON’ state. More intricate models have been proposed to accommodate the complex stochastic dynamics observed experimentally, often relying on a predefined number of gene states and a matrix of rate transitions between them (42, 71). However, the molecular and mechanistic interpretation of these gene states and their transitions is not always evident. In contrast, the SMF approach allows us to propose a stochastic model that explicitly considers the observed promoter states with specific molecular interpretation. For simplicity, we model only the three most important promoter states identified from SMF data: a nucleosome state, where the promoter is closed and occupied by nucleosomes; an open state, consisting of accessible DNA without detectable nucleosomes and Pol II footprints (Unbound and PIC states from SMF); and a Pol II-bound state, where the promoter is occupied by Pol II (PIC + Pol II and Pol II states from SMF). The rate transitions between these states acquire an immediate mechanistic interpretation: the transition between the closed and open states is determined by the effective rates of nucleosome binding and unbinding (*k*_*c*_ and *k*_*o*_), and the transition between the open and Pol II-bound states is determined by the effective Pol II initiation and turnover rates (*k*_*i*_ and *k*_*t*_). Thus, the probabilities of finding the promoter in a given state follow the master equation below:

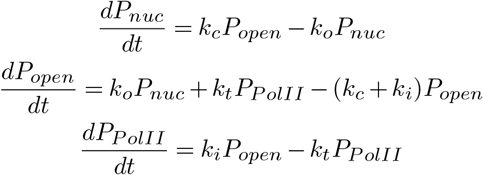

At steady state the probabilities of finding the promoter in an open state or bound by a Pol II molecule are:

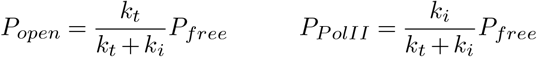

where we define the probability of the promoter being free as *P*_*free*_ = 1 − *P*_*nuc*_. By rearranging the second equation above, we arrive at the main theoretical result of this paper:

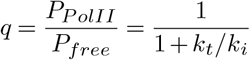

showing that the fraction of promoter molecules occupied by Pol II over the fraction accessible molecules (*q*) follows a simple sigmoid function that depends on the ratio of the turnover rate *k*_*t*_ and the initiation rate *k*_*i*_. In other words, the Pol II occupancy level is determined by the balance of incoming Pol II complexes into the promoter and the Pol II turnover speed.

### Statistical analysis and visualisation

All statistical analyses were carried out using R software (version 4.2.2) (72) and the R package tidyverse (version 2.0.0) (73) and are described in the figure legends and methods section. Plots were generated using ggplot2 (version 3.4.4) (74), ggpubr (version 0.6.0) (75), cowplot (version 1.1.1) (76), and pheatmap (version 1.0.12) (77). The upper and lower boundaries of the box plots represent the 25^th^ and the 75^th^ percentiles. The central line represents the median.

### Data availability

The datasets and computer code produced in this study are available in the following databases:

- Amplicon SMF data: ArrayExpress E-MTAB-14461
- PRO-seq data: ArrayExpress E-MTAB-14462
- Custom scripts for data analyses: https://github.com/Krebslabrep/Chatsirisupachai_Moene_2024

## ACKNOWLEDGEMENTS

We would like to thank members from the Krebs lab, Luca Giorgetti, and László Tora for helpful feedback on the manuscript. Research in the laboratory of AK is supported by core funding from the EMBL, Deutsche Forschungsgemeinschaft (KR 5247/1-2, KR 5247/1-3) and the ERC (TFCoop-101125530). NM developed his work at the Interdisciplinary Thematic Institute IMCBio+, part of the ITI 2021-2028 program of the University of Strasbourg, CNRS and Inserm, supported by IdEx Unistra (ANR-10-IDEX-0002), and by SFRI-STRAT’US project (ANR-20-SFRI-0012) and EUR IMCBio (ANR-17-EURE-0023) under the framework of the France 2030 Program. KC salary is supported by the EMBO Postdoctoral Fellowship (ALTF 482-2022) and DFG (KR 5247/1-3). The authors are grateful to GBCS and GeneCore at EMBL for support in sequencing data generation and management. We thank the Theory@EMBL programme for supporting the sabbatical stay of NM at EMBL.

## AUTHOR CONTRIBUTIONS

KC and AK wrote the manuscript with input from NM and CM. KC generated PRO-seq data with support from RK. CM generated amplicon SMF data with support from RK. KC and CM analysed the data. EK generated SMF data in the somatic cell lines. RK generated amplicon and bait-capture SMF data. NM wrote the mathematical model and integrated the SMF-derived occupancy and PRO-seq data with the help of KC. AK designed and supervised the study. All authors discussed the results and approved the manuscript.

## Supplement Figures

**Fig. S1.**
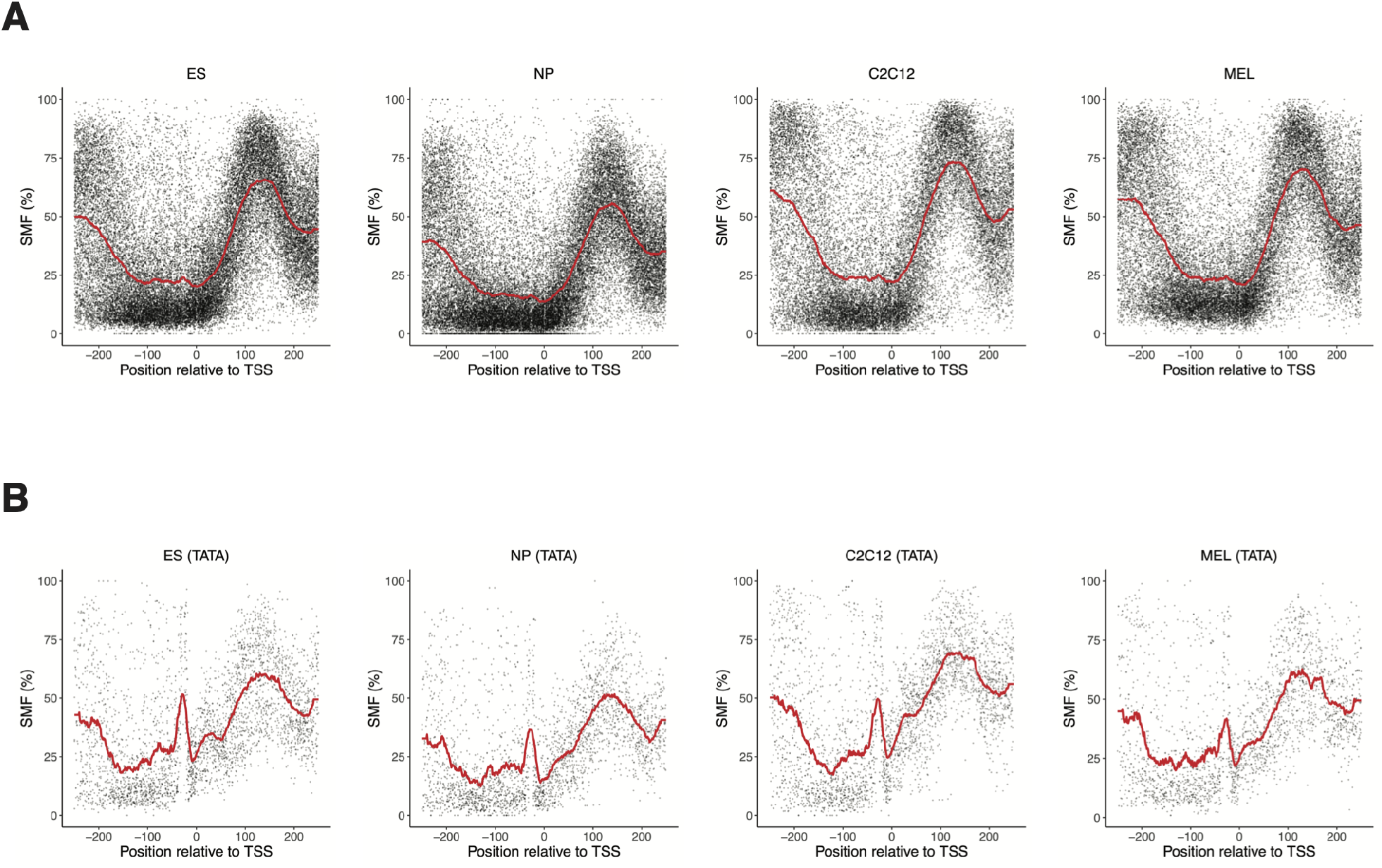
Average promoter footprint of various mouse cell lines. Absence of Pol II footprints at highly active mouse promoters is not only restricted to TKO mESCs but also a property of other mouse cell types. Composite profile of SMF signal at **(A)** the TSSs of active promoters (top 5% Pol II ChIP-seq) and **(B)** the TSSs of active, TATA-box containing promoters (top 5% Pol II ChIP-seq with a TATA-box) in mouse wild-type embryonic stem cells (ES), neural progenitor cells (NP), myoblasts (C2C12), and erythrocytes (MEL).

**Fig. S2.**
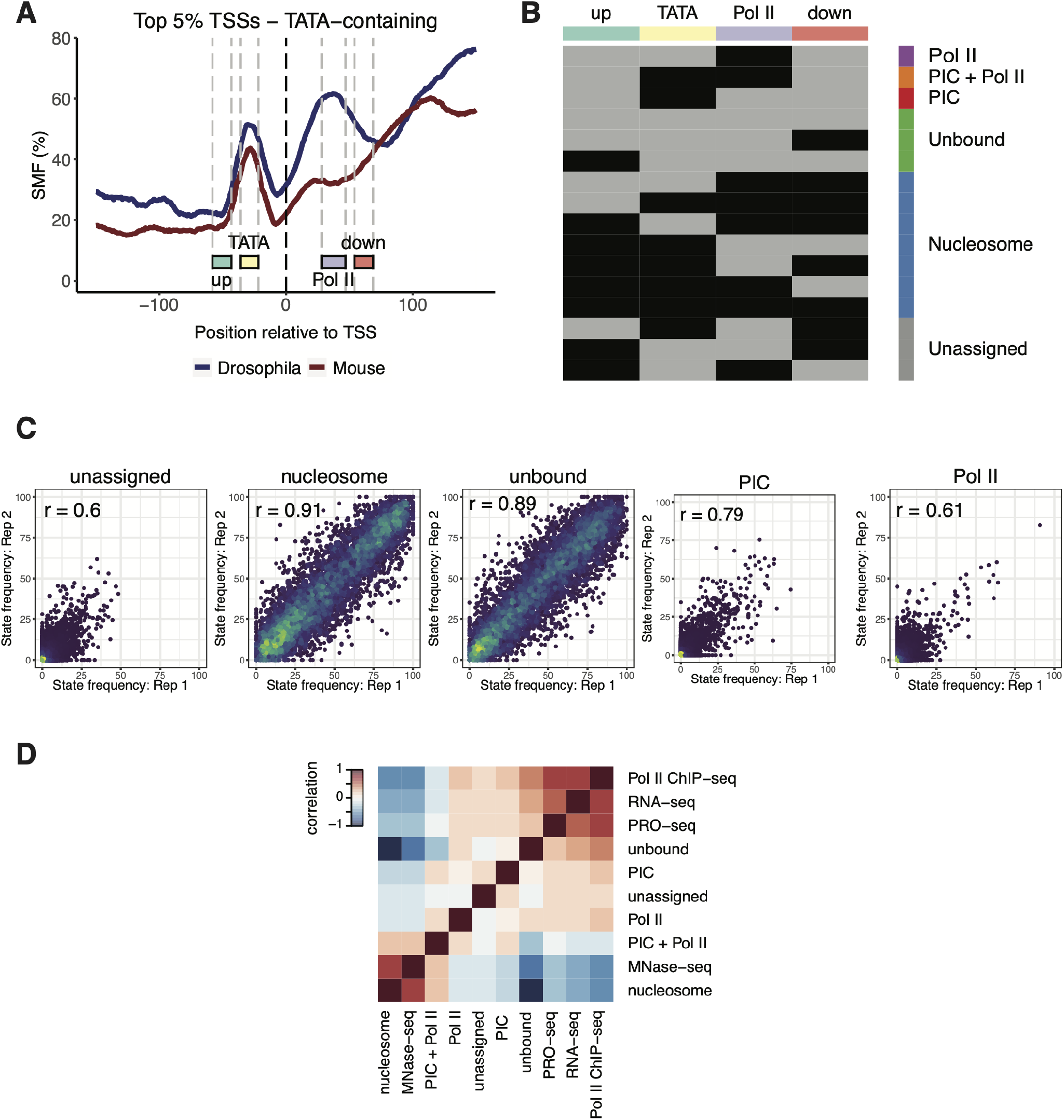
Single-molecule promoter state decomposition in mouse promoters. **(A)** A four-bin strategy to sort single DNA molecule into distinct promoter states. This bin strategy is adapted from (34) to include the downstream bin to better distinguish the Pol II footprint from the nucleosome footprint. Methylation status is binarized within each bin, creating a 4-bit vector that leads to 16 possible combinations, each describing the state of each molecule (see Methods). **(B)** Schematic representation of the methylation patterns used to define each promoter state (methylated Cs – accessible – light grey; unmethylated Cs – protected – black). The top horizontal annotation bar represents the four bins (upstream, TATA, Pol II, downstream) used for promoter state decomposition. The vertical side bar displays the promoter state corresponding to the occupancy type according to the bins. **(C)** Scatter plots show the correlation between state frequencies determined from each bait-capture SMF replicate (n = 6,122 promoters). The states PIC + Pol II and Pol II are combined into a single Pol II state. Pearson correlation coefficients are displayed. **(D)** The global relationship between promoter state frequencies and independent bulk measurements of Pol II and nucleosomes. The heatmap shows the Spearman correlation between each dataset, ordered by hierarchical clustering. The states segregate into two groups correlating either with transcription/transcription machinery (RNA-seq, PRO-seq, and Pol II ChIP-seq) or nucleosomes (MNase-seq).

**Fig. S3.**
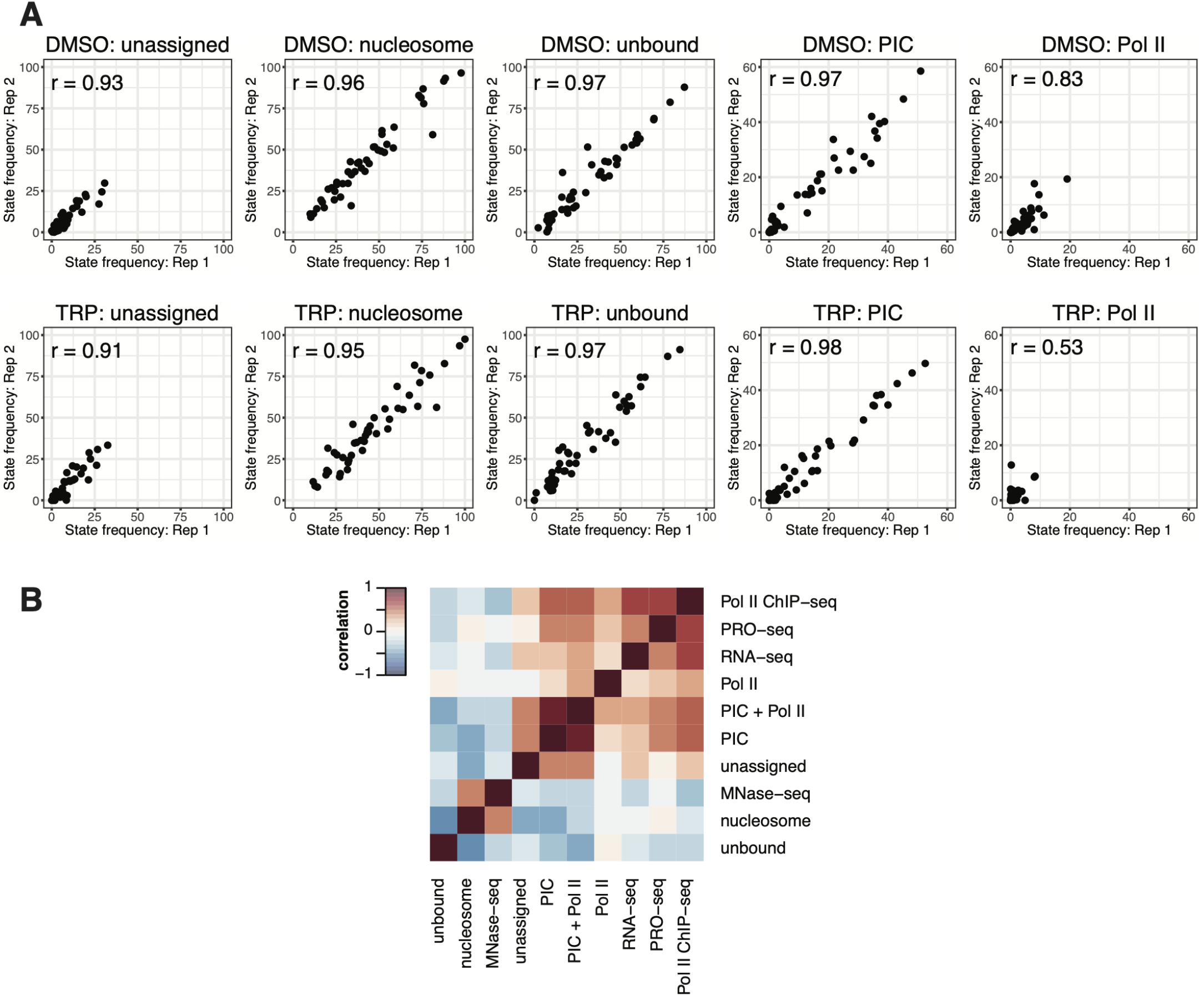
Quality control of targeted amplicon SMF sequencing. **(A)** Scatter plots illustrating the correlation between state frequencies determined from each amplicon SMF replicate. The states PIC + Pol II and Pol II are combined into a single Pol II state. Pearson correlation coefficients are displayed. **(B)** The global relationship between promoter state frequencies at selected promoters that were included in amplicon SMF (n = 47 promoters) and independent bulk measurements of Pol II and nucleosomes. The heatmap shows Spearman correlation between each dataset and is ordered by hierarchical clustering. States separate into two groups which either correlate with transcription/transcription machinery (RNA-seq, PRO-seq, and Pol II ChIP-seq) or nucleosomes (MNase-seq).

**Fig. S4.**
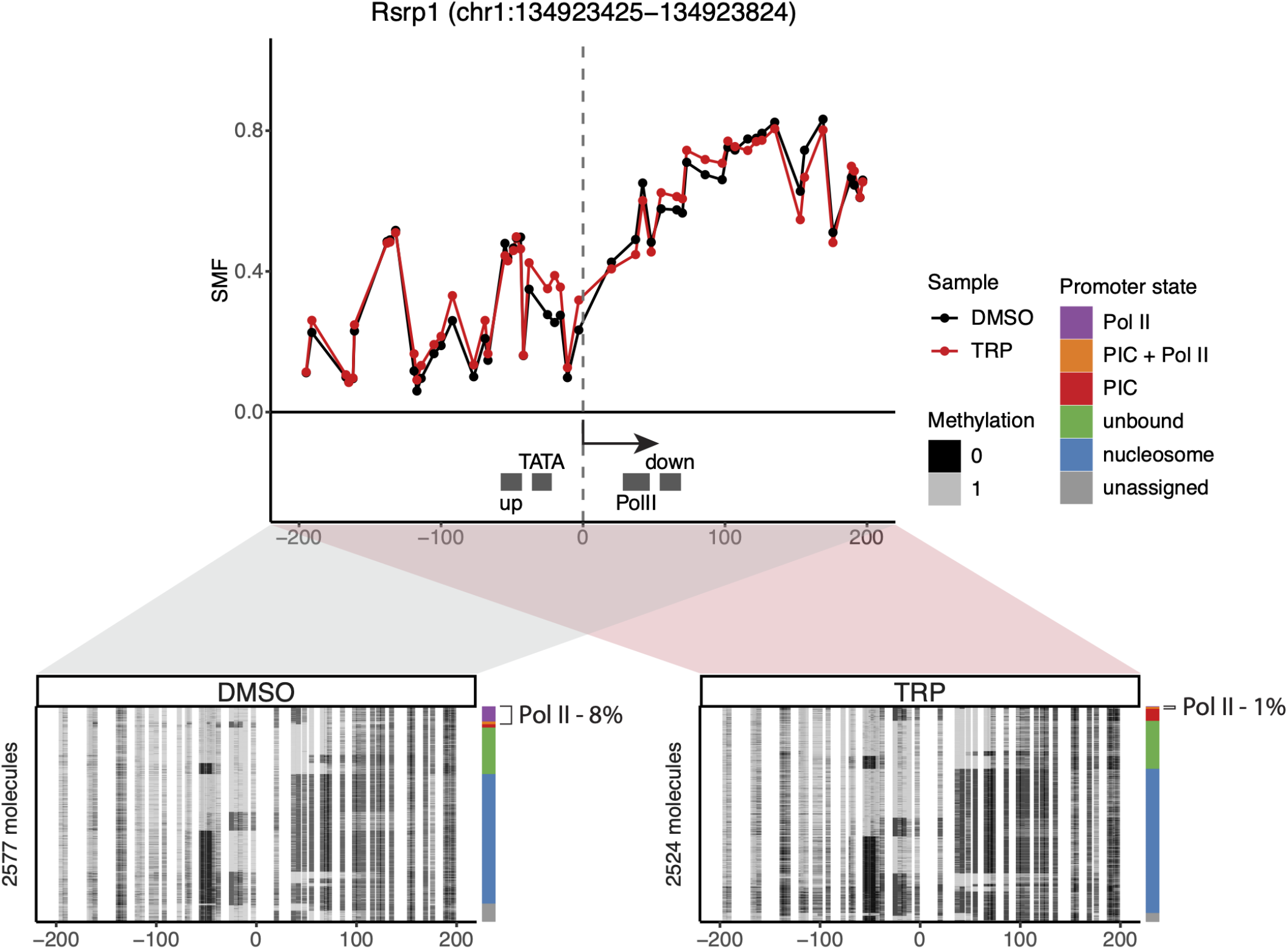
Single-site example of a reduction in Pol II footprint upon TRP treatment at the *Rsrp1* promoter. The upper panel shows the average SMF plot (DMSO – black, TRP – red). The positions of the four bins (upstream, TATA, Pol II, downstream) used for promoter state decomposition are shown (see Methods). The x-axis represents the position relative to the TSS, while the y-axis shows the SMF signal (1 - methylation). The lower panel displays single-molecule stack plots for DMSO and TRP conditions. Each row denotes a single DNA molecule and the methylation status of each cytosine in that molecule (methylated, accessible – light grey; unmethylated, protected – black). The vertical side bars display the frequency of each promoter state. The percentages of molecules harbouring footprints for the engaged Pol II are indicated on the right side of the plot.

**Fig. S5.**
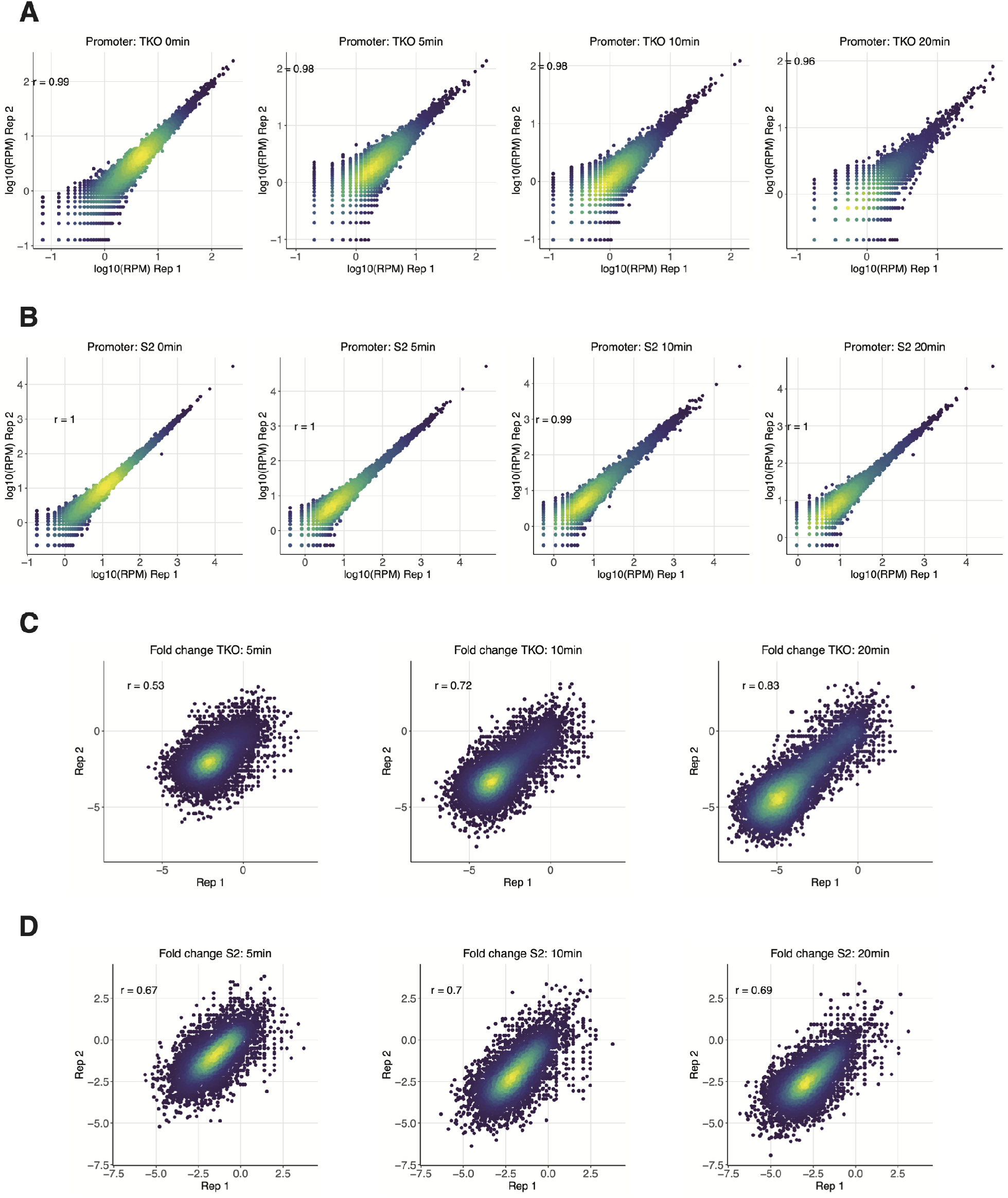
Quality control of PRO-seq data. **(A)** Scatter plots show correlation of the PRO-seq signal (log10(RPM) at the TSS [- 100:200]; n = 24,869 promoters) in mouse TKO mESCs from two replicates of each TRP treatment time point, except in the 2.5 min TRP treatment as only one replicate passed the quality control. Pearson correlation coefficients are displayed. **(B)** Scatter plots show correlation of the PRO-seq signal (log10(RPM) at the TSS [-100:200]; n = 16,198 promoters) in *Drosophila* S2 cells from two replicates of each TRP treatment time point, except in the 2.5 min TRP treatment as only one replicate passed the quality control. Pearson correlation coefficients are displayed. **(C)** Scatter plots show correlation of the log2 fold change in PRO-seq signal upon TRP treatment in mouse TKO mESCs from two replicates, except in the 2.5 min TRP treatment. Pearson correlation coefficients are displayed. **(D)** Scatter plots show correlation of the log2 fold change in PRO-seq signal upon TRP treatment in *Drosophila* S2 cells from two replicates, except in the 2.5 min TRP treatment. Pearson correlation coefficients are displayed.

## Supplement Table

**Table S1.**
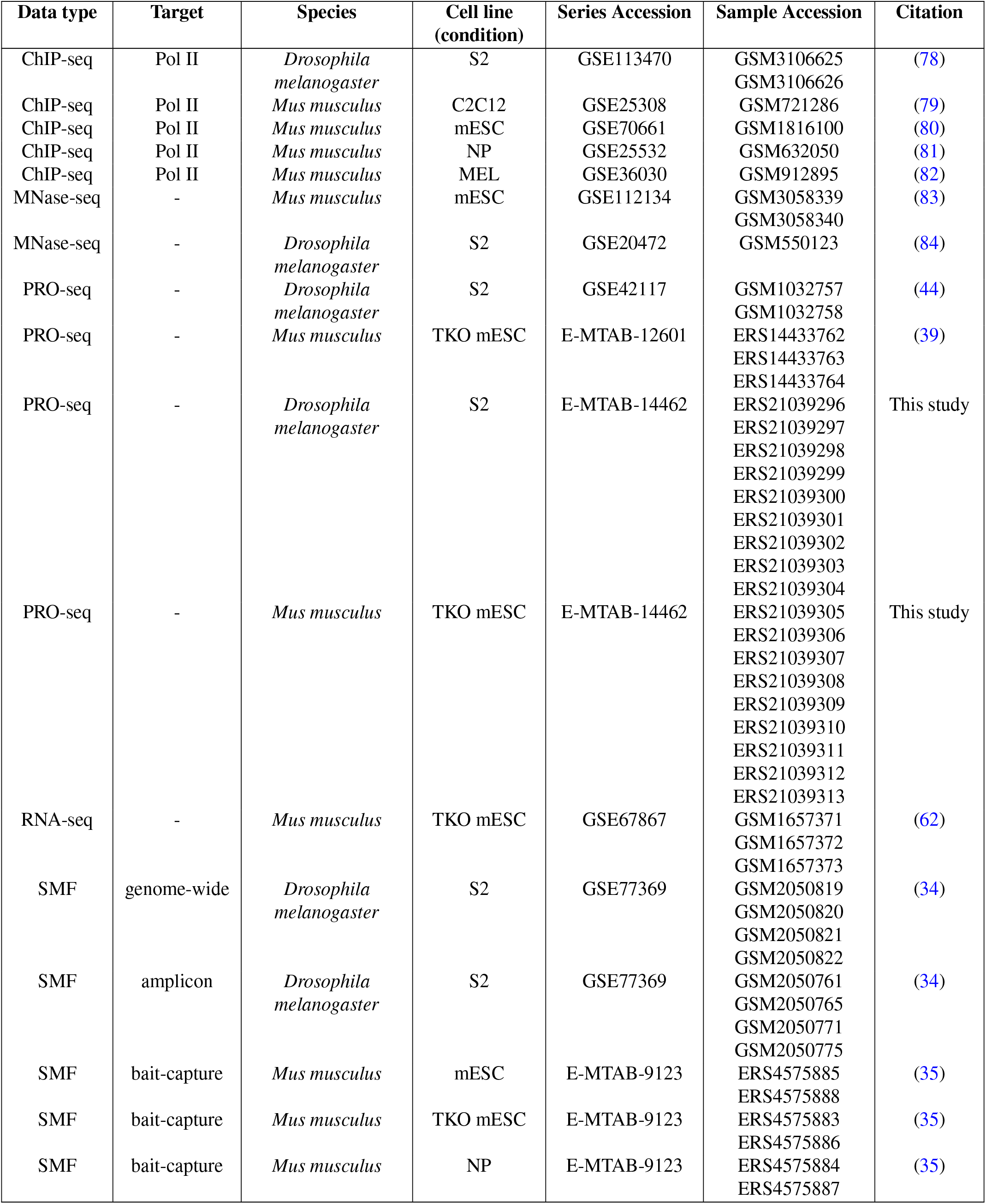
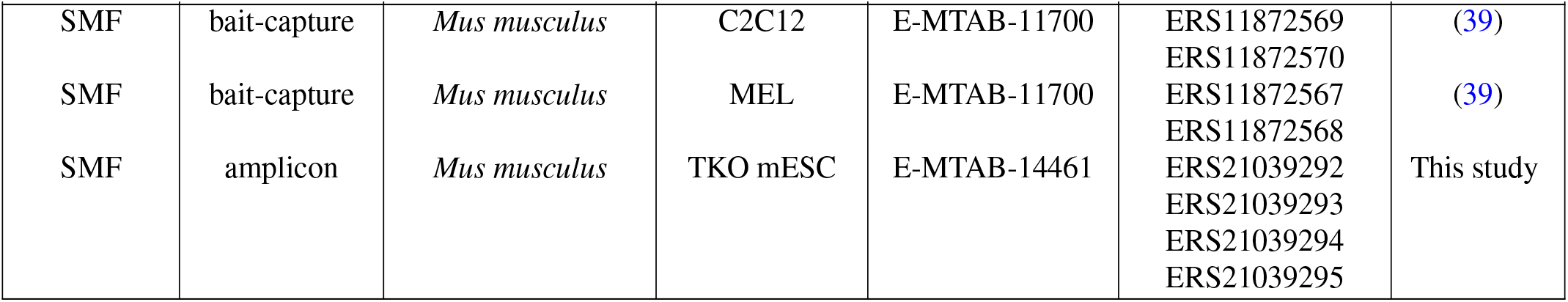
Sequencing datasets used in this study. A list with the details of all sequencing datasets used in this study. For datasets deposited in GEO, the GEO series accession and sample accession are listed, for datasets deposited in ArrayExpress the accession code of the study and the ENA sample code are listed.

## References

1. David L. Bentleys. Multiple forms and functions of premature termination by RNA polymerase II. Journal of Molecular Biology, page 168743, August 2024. ISSN 00222836. doi: 10.1016/j.jmb.2024.168743.

2. Patrick Cramer Organization and regulation of gene transcription. Nature, 573(7772):45–54, September 2019. ISSN 0028-0836, 1476-4687. doi: 10.1038/s41586-019-1517-4.

3. Edward Tunnacliffe and Jonathan R. Chubb. What Is a Transcriptional Burst? Trends in Genetics, 36(4):288–297, April 2020. ISSN 01689525. doi: 10.1016/j.tig.2020.01.003.

4. Anton J. M. Larsson, Per Johnsson, Michael Hagemann-Jensen, Leonard Hartmanis, Omid R. Faridani, Björn Reinius, Åsa Segerstolpe, Chloe M. Rivera, Bing Ren, and Rickard Sandberg Genomic encoding of transcriptional burst kinetics. Nature, 565(7738):251–254, January 2019. ISSN 0028-0836, 1476-4687. doi: 10.1038/s41586-018-0836-1.

5. Hiroshi Ochiai, Tetsutaro Hayashi, Mana Umeda, Mika Yoshimura, Akihito Harada, Yukiko Shimizu, Kenta Nakano, Noriko Saitoh, Zhe Liu, Takashi Yamamoto, Tadashi Okamura, Yasuyuki Ohkawa, Hiroshi Kimura, and Itoshi Nikaido Genome-wide kinetic properties of transcriptional bursting in mouse embryonic stem cells. Science Advances, 6(25):eaaz6699, June 2020. ISSN 2375-2548. doi: 10.1126/sciadv.aaz6699.

6. Daniel Ramsköld, Gert-Jan Hendriks, Anton J. M. Larsson, Juliane V. Mayr, Christoph Ziegenhain, Michael Hagemann-Jensen, Leonard Hartmanis, and Rickard Sandberg Single-cell new RNA sequencing reveals principles of transcription at the resolution of idividual bursts. Nature Cell Biology, August 2024. ISSN 1465-7392, 1476-4679. doi: 10.1038/s41556-024-01486-9.

7. Hernan G. Garcia, Mikhail Tikhonov, Albert Lin, and Thomas Gregor Quantitative Imaging of Transcription in Living Drosophila Embryos Links Polymerase Activity to Patterning. Current Biology, 23(21):2140–2145, November 2013. ISSN 09609822. doi: 10.1016/j.cub.2013.08.054.

8. Kota Hamamoto, Yusuke Umemura, Shiho Makino, and Takashi Fukaya Dynamic interplay between non-coding enhancer transcription and gene activity in development. Nature Communications, 14(1):826, February 2023. ISSN 2041-1723. doi: 10.1038/s41467-023-36485-1.

9. Mounia Lagha, Jacques P. Bothma, Emilia Esposito, Samuel Ng, Laura Stefanik, Chiahao Tsui, Jeffrey Johnston, Kai Chen, David S. Gilmour, Julia Zeitlinger, and Michael S. Levine. Paused Pol II Coordinates Tissue Morphogenesis in the Drosophila Embryo. Cell, 153(5):976–987, May 2013. ISSN 00928674. doi: 10.1016/j.cell.2013.04.045.

10. Virginia L. Pimmett, Matthieu Dejean, Carola Fernandez, Antonio Trullo, Edouard Bertrand, Ovidiu Radulescu, and Mounia Lagha Quantitative imaging of transcription in living Drosophila embryos reveals the impact of core promoter motifs on promoter state dynamics. Nature Communications, 12(1):4504, July 2021. ISSN 2041-1723. doi: 10.1038/s41467-021-24461-6.

11. Joseph Rodriguez, Gang Ren, Christopher R. Day, Keji Zhao, Carson C. Chow, and Daniel R. Larson. Intrinsic Dynamics of a Human Gene Reveal the Basis of Expression Heterogeneity. Cell, 176(1-2):213–226.e18, January 2019. ISSN 00928674. doi: 10.1016/j.cell.2018.11.026.

12. Diana A. Stavreva, David A. Garcia, Gregory Fettweis, Prabhakar R. Gudla, George F. Zaki, Vikas Soni, Andrew McGowan, Geneva Williams, Anh Huynh, Murali Palangat, R. Louis Schiltz, Thomas A. Johnson, Diego M. Presman, Matthew L. Ferguson, Gianluca Pegoraro, Arpita Upadhyaya, and Gordon L. Hager. Transcriptional Bursting and Co-bursting Regulation by Steroid Hormone Release Pattern and Transcription Factor Mobility. Molecular Cell, 75(6):1161–1177.e11, September 2019. ISSN 10972765. doi: 10.1016/j.molcel.2019.06.042.

13. David M. Suter, Nacho Molina, David Gatfield, Kim Schneider, Ueli Schibler, and Felix Naef. Mammalian Genes Are Transcribed with Widely Different Bursting Kinetics. Science, 332 (6028):472–474, April 2011. ISSN 0036-8075, 1095-9203. doi: 10.1126/science.1198817.

14. Yihan Wan, Dimitrios G. Anastasakis, Joseph Rodriguez, Murali Palangat, Prabhakar Gudla, George Zaki, Mayank Tandon, Gianluca Pegoraro, Carson C. Chow, Markus Hafner, and Daniel R. Larson. Dynamic imaging of nascent RNA reveals general principles of transcription dynamics and stochastic splice site selection. Cell, 184(11):2878–2895.e20, May 2021. ISSN 00928674. doi: 10.1016/j.cell.2021.04.012.

15. Nicholas C. Lammers, Yang Joon Kim, Jiaxi Zhao, and Hernan G. Garcia. A matter of time: Using dynamics and theory to uncover mechanisms of transcriptional bursting. Current Opinion in Cell Biology, 67:147–157, December 2020. ISSN 09550674. doi: 10.1016/j.ceb.2020.08.001.

16. Joseph V.W. Meeussen and Tineke L. Lenstra. Time will tell: comparing timescales to gain insight into transcriptional bursting. Trends in Genetics, page S0168952523002585, January 2024. ISSN 01689525. doi: 10.1016/j.tig.2023.11.003.

17. R Abdella, A Talyzina, S Chen, CJ Inouye, R Tjian, and Y He. Structure of the human Mediator-bound transcription preinitiation complex. 2021.

18. Shintaro Aibara, Sandra Schilbach, and Patrick Cramer. Structures of mammalian RNA polymerase II pre-initiation complexes. Nature, 594(7861):124–128, June 2021. ISSN 0028-0836, 1476-4687. doi: 10.1038/s41586-021-03554-8.

19. Merle Hantsche and Patrick Cramer. Conserved RNA polymerase II initiation complex structure. Current Opinion in Structural Biology, 47:17–22, December 2017. ISSN 0959440X. doi: 10.1016/j.sbi.2017.03.013.

20. Yuan He, Jie Fang, Dylan J. Taatjes, and Eva Nogales. Structural visualization of key steps in human transcription initiation. Nature, 495(7442):481–486, March 2013. ISSN 0028-0836, 1476-4687. doi: 10.1038/nature11991.

21. Wolfgang Mühlbacher, Sarah Sainsbury, Matthias Hemann, Merle Hantsche, Simon Neyer, Franz Herzog, and Patrick Cramer. Conserved architecture of the core RNA polymerase II initiation complex. Nature Communications, 5(1):4310, July 2014. ISSN 2041-1723. doi: 10.1038/ncomms5310.

22. Eva Nogales, Robert K. Louder, and Yuan He. Structural Insights into the Eukaryotic Transcription Initiation Machinery. Annual Review of Biophysics, 46(1):59–83, May 2017. ISSN 1936-122X, 1936-1238. doi: 10.1146/annurev-biophys-070816-033751.

23. Alexandra G. Chivu, Abderhman Abuhashem, Gilad Barshad, Edward J. Rice, Michelle M. Leger, Albert C. Vill, Wilfred Wong, Rebecca Brady, Jeramiah J. Smith, Athula H. Wikramanayake, César Arenas-Mena, Ilana L. Brito, Iñaki Ruiz-Trillo, Anna-Katerina Hadjantonakis, John T. Lis, James J. Lewis, and Charles G. Danko. Evolution of promoter-proximal pausing enabled a new layer of transcription control. bioRxiv, 2023. doi: 10.1101/2023.02.19.529146. Publisher: Cold Spring Harbor Laboratory _eprint: https://www.biorxiv.org/content/early/2023/02/19/2023.02.19.529146.full.pdf.

24. Chad B. Stein, Andrew R. Field, Claudia A. Mimoso, ChenCheng Zhao, Kai-Lieh Huang, Eric J. Wagner, and Karen Adelman. Integrator endonuclease drives promoter-proximal termination at all RNA polymerase II-transcribed loci. Molecular Cell, 82(22):4232–4245.e11, November 2022. ISSN 10972765. doi: 10.1016/j.molcel.2022.10.004.

25. Seychelle M Vos, David Pöllmann, Livia Caizzi, Katharina B Hofmann, Pascaline Rombaut, Tomasz Zimniak, Franz Herzog, and Patrick Cramer. Architecture and RNA binding of the human negative elongation factor. eLife, 5:e14981, June 2016. ISSN 2050-084X. doi: 10.7554/eLife.14981.

26. Sarah A. Welsh and Alessandro Gardini. Genomic regulation of transcription and RNA processing by the multitasking Integrator complex. Nature Reviews Molecular Cell Biology, 24 (3):204–220, March 2023. ISSN 1471-0072, 1471-0080. doi: 10.1038/s41580-022-00534-2.

27. Telmo Henriques, Daniel A. Gilchrist, Sergei Nechaev, Michael Bern, Ginger W. Muse, Adam Burkholder, David C. Fargo, and Karen Adelman. Stable Pausing by RNA Polymerase II Provides an Opportunity to Target and Integrate Regulatory Signals. Molecular Cell, 52(4):517–528, November 2013. ISSN 10972765. doi: 10.1016/j.molcel.2013.10.001.

28. Lucy H. Williams, George Fromm, Nolan G. Gokey, Telmo Henriques, Ginger W. Muse, Adam Burkholder, David C. Fargo, Guang Hu, and Karen Adelman. Pausing of RNA Polymerase II Regulates Mammalian Developmental Potential through Control of Signaling Networks. Molecular Cell, 58(2):311–322, April 2015. ISSN 10972765. doi: 10.1016/j.molcel.2015.02.003.

29. Qiye He, Jeff Johnston, and Julia Zeitlinger. ChIP-nexus enables improved detection of in vivo transcription factor binding footprints. Nature Biotechnology, 33(4):395–401, April 2015. ISSN 1087-0156, 1546-1696. doi: 10.1038/nbt.3121.

30. Ho Sung Rhee and B. Franklin Pugh. Comprehensive Genome-wide Protein-DNA Interactions Detected at Single-Nucleotide Resolution. Cell, 147(6):1408–1419, December 2011. ISSN 00928674. doi: 10.1016/j.cell.2011.11.013.

31. Matthew J. Rossi, Prashant K. Kuntala, William K. M. Lai, Naomi Yamada, Nitika Badjatia, Chitvan Mittal, Guray Kuzu, Kylie Bocklund, Nina P. Farrell, Thomas R. Blanda, Joshua D. Mairose, Ann V. Basting, Katelyn S. Mistretta, David J. Rocco, Emily S. Perkinson, Gretta D. Kellogg, Shaun Mahony, and B. Franklin Pugh. A high-resolution protein architecture of the budding yeast genome. Nature, 592(7853):309–314, April 2021. ISSN 0028-0836, 1476-4687. doi: 10.1038/s41586-021-03314-8.

32. Wanqing Shao and Julia Zeitlinger. Paused RNA polymerase II inhibits new transcriptional initiation. Nature Genetics, 49(7):1045–1051, July 2017. ISSN 1061-4036, 1546-1718. doi: 10.1038/ng.3867.

33. Rozemarijn W. D. Kleinendorst, Guido Barzaghi, Mike L. Smith, Judith B. Zaugg, and Arnaud R. Krebs. Genome-wide quantification of transcription factor binding at single-DNA-molecule resolution using methyl-transferase footprinting. Nature Protocols, 16(12):5673–5706, December 2021. ISSN 1754-2189, 1750-2799. doi: 10.1038/s41596-021-00630-1.

34. Arnaud R. Krebs, Dilek Imanci, Leslie Hoerner, Dimos Gaidatzis, Lukas Burger, and Dirk Schübeler. Genome-wide Single-Molecule Footprinting Reveals High RNA Polymerase II Turnover at Paused Promoters. Molecular Cell, 67(3):411–422.e4, August 2017. ISSN 10972765. doi: 10.1016/j.molcel.2017.06.027.

35. Can Sönmezer, Rozemarijn Kleinendorst, Dilek Imanci, Guido Barzaghi, Laura Villacorta, Dirk Schübeler, Vladimir Benes, Nacho Molina, and Arnaud Regis Krebs. Molecular Co-occupancy Identifies Transcription Factor Binding Cooperativity In Vivo. Molecular Cell, 81 (2):255–267.e6, January 2021. ISSN 10972765. doi: 10.1016/j.molcel.2020.11.015.

36. Imad Abugessaisa, Shuhei Noguchi, Akira Hasegawa, Jayson Harshbarger, Atsushi Kondo, Marina Lizio, Jessica Severin, Piero Carninci, Hideya Kawaji, and Takeya Kasukawa. FANTOM5 CAGE profiles of human and mouse reprocessed for GRCh38 and GRCm38 genome assemblies. Scientific Data, 4(1):170107, August 2017. ISSN 2052-4463. doi: 10.1038/sdata.2017.107.

37. Thomas W. Tullius, R. Stefan Isaac, Danilo Dubocanin, Jane Ranchalis, L. Stirling Churchman, and Andrew B. Stergachis. RNA polymerases reshape chromatin architecture and couple transcription on individual fibers. Molecular Cell, page S1097276524006671, August 2024. ISSN 10972765. doi: 10.1016/j.molcel.2024.08.013.

38. Travis N. Mavrich, Cizhong Jiang, Ilya P. Ioshikhes, Xiaoyong Li, Bryan J. Venters, Sara J. Zanton, Lynn P. Tomsho, Ji Qi, Robert L. Glaser, Stephan C. Schuster, David S. Gilmour, Istvan Albert, and B. Franklin Pugh. Nucleosome organization in the Drosophila genome. Nature, 453(7193):358–362, May 2008. ISSN 0028-0836, 1476-4687. doi: 10.1038/nature06929.

39. Elisa Kreibich, Rozemarijn Kleinendorst, Guido Barzaghi, Sarah Kaspar, and Arnaud R. Krebs. Single-molecule footprinting identifies context-dependent regulation of enhancers by DNA methylation. Molecular Cell, 83(5):787–802.e9, March 2023. ISSN 10972765. doi: 10.1016/j.molcel.2023.01.017.

40. Iris Jonkers, Hojoong Kwak, and John T Lis. Genome-wide dynamics of Pol II elongation and its interplay with promoter proximal pausing, chromatin, and exons. eLife, 3:e02407, April 2014. ISSN 2050-084X. doi: 10.7554/eLife.02407.

41. J. Peccoud and B. Ycart. Markovian Modeling of Gene-Product Synthesis. Theoretical Population Biology, 48(2):222–234, October 1995. ISSN 00405809. doi: 10.1006/tpbi.1995.1027.

42. Benjamin Zoller, Damien Nicolas, Nacho Molina, and Felix Naef. Structure of silent transcription intervals and noise characteristics of mammalian genes. Molecular Systems Biology, 11(7):823, July 2015. ISSN 1744-4292, 1744-4292. doi: 10.15252/msb.20156257.

43. Julius Judd, Luke A. Wojenski, Lauren M. Wainman, Nathaniel D. Tippens, Edward J. Rice, Alexis Dziubek, Geno J. Villafano, Erin M. Wissink, Philip Versluis, Lina Bagepalli, Sagar R. Shah, Dig B. Mahat, Jacob M. Tome, Charles G. Danko, John T. Lis, and Leighton J. Core. A rapid, sensitive, scalable method for Precision Run-On sequencing (PRO-seq), May 2020.

44. Hojoong Kwak, Nicholas J. Fuda, Leighton J. Core, and John T. Lis. Precise Maps of RNA Polymerase Reveal How Promoters Direct Initiation and Pausing. Science, 339(6122):950–953, February 2013. ISSN 0036-8075, 1095-9203. doi: 10.1126/science.1229386.

45. Sergei Nechaev, David C. Fargo, Gilberto Dos Santos, Liwen Liu, Yuan Gao, and Karen Adelman. Global Analysis of Short RNAs Reveals Widespread Promoter-Proximal Stalling and Arrest of Pol II in Drosophila. Science, 327(5963):335–338, January 2010. ISSN 0036-8075, 1095-9203. doi: 10.1126/science.1181421.

46. Arnaud R. Krebs. Studying transcription factor function in the genome at molecular resolution. Trends in Genetics, 37(9):798–806, September 2021. ISSN 01689525. doi: 10.1016/j.tig.2021.03.008.

47. Leighton Core and Karen Adelman. Promoter-proximal pausing of RNA polymerase II: a nexus of gene regulation. Genes & Development, 33(15-16):960–982, August 2019. ISSN 0890-9369, 1549-5477. doi: 10.1101/gad.325142.119.

48. Nicholas J. Fuda, M. Behfar Ardehali, and John T. Lis. Defining mechanisms that regulate RNA polymerase II transcription in vivo. Nature, 461(7261):186–192, September 2009. ISSN 0028-0836, 1476-4687. doi: 10.1038/nature08449.

49. Karen Adelman and John T. Lis. Promoter-proximal pausing of RNA polymerase II: emerging roles in metazoans. Nature Reviews Genetics, 13(10):720–731, October 2012. ISSN 1471-0056, 1471-0064. doi: 10.1038/nrg3293.

50. Xavier Darzacq, Yaron Shav-Tal, Valeria De Turris, Yehuda Brody, Shailesh M Shenoy, Robert D Phair, and Robert H Singer. In vivo dynamics of RNA polymerase II transcription. Nature Structural & Molecular Biology, 14(9):796–806, September 2007. ISSN 1545-9993, 1545-9985. doi: 10.1038/nsmb1280.

51. Benjamin Erickson, Ryan M. Sheridan, Michael Cortazar, and David L. Bentley. Dynamic turnover of paused Pol II complexes at human promoters. Genes & Development, 32(17-18): 1215–1225, September 2018. ISSN 0890-9369, 1549-5477. doi: 10.1101/gad.316810.118.

52. Kyle A. Nilson, Christine K. Lawson, Nicholas J. Mullen, Christopher B. Ball, Benjamin M. Spector, Jeffery L. Meier, and David H. Price. Oxidative stress rapidly stabilizes promoter-proximal paused Pol II across the human genome. Nucleic Acids Research, 45(19):11088–11105, November 2017. ISSN 0305-1048, 1362-4962. doi: 10.1093/nar/gkx724.

53. David H. Price. Transient pausing by RNA polymerase II. Proceedings of the National Academy of Sciences, 115(19):4810–4812, May 2018. ISSN 0027-8424, 1091-6490. doi: 10.1073/pnas.1805129115.

54. Barbara Steurer, Roel C. Janssens, Bart Geverts, Marit E. Geijer, Franziska Wienholz, Arjan F. Theil, Jiang Chang, Shannon Dealy, Joris Pothof, Wiggert A. Van Cappellen, Adriaan B. Houtsmuller, and Jurgen A. Marteijn. Live-cell analysis of endogenous GFP-RPB1 uncovers rapid turnover of initiating and promoter-paused RNA Polymerase II. Proceedings of the National Academy of Sciences, 115(19), May 2018. ISSN 0027-8424, 1091-6490. doi: 10.1073/pnas.1717920115.

55. Nathan D. Elrod, Telmo Henriques, Kai-Lieh Huang, Deirdre C. Tatomer, Jeremy E. Wilusz, Eric J. Wagner, and Karen Adelman. The Integrator Complex Attenuates Promoter-Proximal Transcription at Protein-Coding Genes. Molecular Cell, 76(5):738–752.e7, December 2019. ISSN 10972765. doi: 10.1016/j.molcel.2019.10.034.

56. Isaac Fianu, Moritz Ochmann, James L. Walshe, Olexandr Dybkov, Joseph Neos Cruz, Henning Urlaub, and Patrick Cramer. Structural basis of Integrator-dependent RNA polymerase II termination. Nature, 629(8010):219–227, May 2024. ISSN 0028-0836, 1476-4687. doi: 10.1038/s41586-024-07269-4.

57. Michal Razew, Angelique Fraudeau, Moritz M. Pfleiderer, Romain Linares, and Wojciech P. Galej. Structural basis of the Integrator complex assembly and association with transcription factors. Molecular Cell, 84(13):2542–2552.e5, July 2024. ISSN 10972765. doi: 10.1016/j.molcel.2024.05.009.

58. Eric J. Wagner, Liang Tong, and Karen Adelman. Integrator is a global promoter-proximal termination complex. Molecular Cell, 83(3):416–427, February 2023. ISSN 10972765. doi: 10.1016/j.molcel.2022.11.012.

59. Arjun Raj, Charles S Peskin, Daniel Tranchina, Diana Y Vargas, and Sanjay Tyagi. Stochastic mRNA Synthesis in Mammalian Cells. PLoS Biology, 4(10):e309, September 2006. ISSN 1545-7885. doi: 10.1371/journal.pbio.0040309.

60. Katjana Tantale, Encar Garcia-Oliver, Marie-Cécile Robert, Adèle L’Hostis, Yueyuxiao Yang, Nikolay Tsanov, Rachel Topno, Thierry Gostan, Alja Kozulic-Pirher, Meenakshi Basu-Shrivastava, Kamalika Mukherjee, Vera Slaninova, Jean-Christophe Andrau, Florian Mueller, Eugenia Basyuk, Ovidiu Radulescu, and Edouard Bertrand. Stochastic pausing at latent HIV-1 promoters generates transcriptional bursting. Nature Communications, 12(1):4503, July 2021. ISSN 2041-1723. doi: 10.1038/s41467-021-24462-5.

61. Benjamin R. Doughty, Michaela M. Hinks, Julia M. Schaepe, Georgi K. Marinov, Abby R. Thurm, Carolina Rios-Martinez, Benjamin E. Parks, Yingxuan Tan, Emil Marklund, Danilo Dubocanin, Lacramioara Bintu, and William J. Greenleaf. Single-molecule chromatin configurations link transcription factor binding to expression in human cells. bioRxiv, 2024. doi: 10.1101/2024.02.02.578660. Publisher: Cold Spring Harbor Laboratory _eprint: https://www.biorxiv.org/content/early/2024/02/04/2024.02.02.578660.full.pdf.

62. Silvia Domcke, Anaïs Flore Bardet, Paul Adrian Ginno, Dominik Hartl, Lukas Burger, and Dirk Schübeler. Competition between DNA methylation and transcription factors determines binding of NRF1. Nature, 528(7583):575–579, December 2015. ISSN 0028-0836, 1476-4687. doi: 10.1038/nature16462.

63. F. Krueger, F. James, P. Ewels, E. Afyounian, and B. Schuster-Boeckler. FelixKrueger/TrimGalore, v0.6.7 - via Zenodo., 2021.

64. Dimos Gaidatzis, Anita Lerch, Florian Hahne, and Michael B. Stadler. QuasR: quantification and annotation of short reads in R. Bioinformatics, 31(7):1130–1132, April 2015. ISSN 1367-4811, 1367-4803. doi: 10.1093/bioinformatics/btu781.

65. Ben Langmead, Cole Trapnell, Mihai Pop, and Steven L Salzberg. Ultrafast and memory-efficient alignment of short DNA sequences to the human genome. Genome Biology, 10(3):R25, 2009. ISSN 1465-6906. doi: 10.1186/gb-2009-10-3-r25.

66. Broad Institute. Picard Toolkit, Broad Institute, GitHub Repository, 2019.

67. Marcel Martin. Cutadapt removes adapter sequences from high-throughput sequencing reads.EMBnet.journal; Vol 17, No 1: Next Generation Sequencing Data Analysis DO-10.14806/ej.17.1.200, May 2011.

68. Tom Smith, Andreas Heger, and Ian Sudbery. UMI-tools: modeling sequencing errors in Unique Molecular Identifiers to improve quantification accuracy. Genome Research, 27(3):491–499, March 2017. ISSN 1088-9051, 1549-5469. doi: 10.1101/gr.209601.116.

69. The FANTOM Consortium and the RIKEN PMI and CLST (DGT). A promoter-level mammalian expression atlas. Nature, 507(7493):462–470, March 2014. ISSN 0028-0836, 1476-4687. doi: 10.1038/nature13182.

70. The modENCODE Consortium, Sushmita Roy, Jason Ernst, Peter V. Kharchenko, Pouya Kheradpour, Nicolas Negre, Matthew L. Eaton, Jane M. Landolin, Christopher A. Bristow, Lijia Ma, Michael F. Lin, Stefan Washietl, Bradley I. Arshinoff, Ferhat Ay, Patrick E. Meyer, Nicolas Robine, Nicole L. Washington, Luisa Di Stefano, Eugene Berezikov, Christopher D. Brown, Rogerio Candeias, Joseph W. Carlson, Adrian Carr, Irwin Jungreis, Daniel Marbach, Rachel Sealfon, Michael Y. Tolstorukov, Sebastian Will, Artyom A. Alekseyenko, Carlo Artieri, Benjamin W. Booth, Angela N. Brooks, Qi Dai, Carrie A. Davis, Michael O. Duff, Xin Feng, Andrey A. Gorchakov, Tingting Gu, Jorja G. Henikoff, Philipp Kapranov, Renhua Li, Heather K. MacAlpine, John Malone, Aki Minoda, Jared Nordman, Katsutomo Okamura, Marc Perry, Sara K. Powell, Nicole C. Riddle, Akiko Sakai, Anastasia Samsonova, Jeremy E. Sandler, Yuri B. Schwartz, Noa Sher, Rebecca Spokony, David Sturgill, Marijke Van Baren, Kenneth H. Wan, Li Yang, Charles Yu, Elise Feingold, Peter Good, Mark Guyer, Rebecca Lowdon, Kami Ahmad, Justen Andrews, Bonnie Berger, Steven E. Brenner, Michael R. Brent, Lucy Cherbas, Sarah C. R. Elgin, Thomas R. Gingeras, Robert Grossman, Roger A. Hoskins, Thomas C. Kaufman, William Kent, Mitzi I. Kuroda, Terry Orr-Weaver, Norbert Perrimon, Vincenzo Pirrotta, James W. Posakony, Bing Ren, Steven Russell, Peter Cherbas, Brenton R. Graveley, Suzanna Lewis, Gos Micklem, Brian Oliver, Peter J. Park, Susan E. Celniker, Steven Henikoff, Gary H. Karpen, Eric C. Lai, David M. MacAlpine, Lincoln D. Stein, Kevin P. White, Manolis Kellis, David Acevedo, Richard Auburn, Galt Barber, Hugo J. Bellen, Eric P. Bishop, Terri D. Bryson, Aurelien Chateigner, Jia Chen, Hiram Clawson, Charles L. G. Comstock, Sergio Contrino, Leyna C. DeNapoli, Queying Ding, Alex Dobin, Marc H. Domanus, Jorg Drenkow, Sandrine Dudoit, Jackie Dumais, Thomas Eng, Delphine Fagegaltier, Sarah E. Gadel, Srinka Ghosh, Francois Guillier, David Hanley, Gregory J. Hannon, Kasper D. Hansen, Elizabeth Heinz, Angie S. Hinrichs, Martin Hirst, Sonali Jha, Lichun Jiang, Youngsook L. Jung, Helena Kashevsky, Cameron D. Kennedy, Ellen T. Kephart, Laura Langton, Ok-Kyung Lee, Sharon Li, Zirong Li, Wei Lin, Daniela Linder-Basso, Paul Lloyd, Rachel Lyne, Sarah E. Marchetti, Marco Marra, Nicolas R. Mattiuzzo, Sheldon McKay, Folker Meyer, David Miller, Steven W. Miller, Richard A. Moore, Carolyn A. Morrison, Joseph A. Prinz, Michelle Rooks, Richard Moore, Kim M. Rutherford, Peter Ruzanov, Douglas A. Scheftner, Lionel Senderowicz, Parantu K. Shah, Gregory Shanower, Richard Smith, E. O. Stinson, Sarah Suchy, Aaron E. Tenney, Feng Tian, Koen J. T. Venken, Huaien Wang, Robert White, Jared Wilkening, Aarron T. Willingham, Chris Zaleski, Zheng Zha, Dayu Zhang, Yongjun Zhao, and Jennifer Zieba. Identification of Functional Elements and Regulatory Circuits by Drosophila modENCODE. Science, 330(6012):1787–1797, December 2010. ISSN 0036-8075, 1095-9203. doi: 10.1126/science.1198374.

71. Katjana Tantale, Florian Mueller, Alja Kozulic-Pirher, Annick Lesne, Jean-Marc Victor, Marie-Cécile Robert, Serena Capozi, Racha Chouaib, Volker Bäcker, Julio Mateos-Langerak, Xavier Darzacq, Christophe Zimmer, Eugenia Basyuk, and Edouard Bertrand. A single-molecule view of transcription reveals convoys of RNA polymerases and multi-scale bursting. Nature Communications, 7(1):12248, July 2016. ISSN 2041-1723. doi: 10.1038/ncomms12248.

72. R Core Team. R: A language and environment for statistical computing., 2022.

73. Hadley Wickham, Mara Averick, Jennifer Bryan, Winston Chang, Lucy McGowan, Romain François, Garrett Grolemund, Alex Hayes, Lionel Henry, Jim Hester, Max Kuhn, Thomas Pedersen, Evan Miller, Stephan Bache, Kirill Müller, Jeroen Ooms, David Robinson, Dana Seidel, Vitalie Spinu, Kohske Takahashi, Davis Vaughan, Claus Wilke, Kara Woo, and Hiroaki Yutani. Welcome to the Tidyverse. Journal of Open Source Software, 4(43):1686, November 2019. ISSN 2475-9066. doi: 10.21105/joss.01686.

74. Hadley Wickham. ggplot2: Elegant Graphics for Data Analysis. Springer-Verlag New York, 2016. ISBN 978-3-319-24277-4.

75. Alboukadel Kassambara. ggpubr: ‘ggplot2’ Based Publication Ready Plots, 2023.

76. Claus O. Wilke. cowplot: Streamlined Plot Theme and Plot Annotations for ‘ggplot2’, 2020.

77. Raivo Kolde. pheatmap: Pretty Heatmaps, 2019.

78. Theophilus T. Tettey, Xin Gao, Wanqing Shao, Hua Li, Benjamin A. Story, Alex D. Chitsazan, Robert L. Glaser, Zach H. Goode, Christopher W. Seidel, Ronald C. Conaway, Julia Zeitlinger, Marco Blanchette, and Joan W. Conaway. A Role for FACT in RNA Polymerase II Promoter-Proximal Pausing. Cell Reports, 27(13):3770–3779.e7, June 2019. ISSN 22111247. doi: 10.1016/j.celrep.2019.05.099.

79. Patrik Asp, Roy Blum, Vasupradha Vethantham, Fabio Parisi, Mariann Micsinai, Jemmie Cheng, Christopher Bowman, Yuval Kluger, and Brian David Dynlacht. Genome-wide remodeling of the epigenetic landscape during myogenic differentiation. Proceedings of the National Academy of Sciences, 108(22), May 2011. ISSN 0027-8424, 1091-6490. doi: 10.1073/pnas.1102223108.

80. Diana Langer, Igor Martianov, Daniel Alpern, Muriel Rhinn, Céline Keime, Pascal Dollé, Gabrielle Mengus, and Irwin Davidson. Essential role of the TFIID subunit TAF4 in murine embryogenesis and embryonic stem cell differentiation. Nature Communications, 7(1):11063, March 2016. ISSN 2041-1723. doi: 10.1038/ncomms11063.

81. Vijay K Tiwari, Michael B Stadler, Christiane Wirbelauer, Renato Paro, Dirk Schübeler, and Christian Beisel. A chromatin-modifying function of JNK during stem cell differentiation. Nature Genetics, 44(1):94–100, January 2012. ISSN 1061-4036, 1546-1718. doi: 10.1038/ng.1036.

82. The ENCODE Project Consortium. An integrated encyclopedia of DNA elements in the human genome. Nature, 489(7414):57–74, September 2012. ISSN 0028-0836, 1476-4687. doi: 10.1038/nature11247.

83. Darko Barisic, Michael B. Stadler, Mario Iurlaro, and Dirk Schübeler. Mammalian ISWI and SWI/SNF selectively mediate binding of distinct transcription factors. Nature, 569(7754):136–140, May 2019. ISSN 0028-0836, 1476-4687. doi: 10.1038/s41586-019-1115-5.

84. Daniel A. Gilchrist, Gilberto Dos Santos, David C. Fargo, Bin Xie, Yuan Gao, Leping Li, and Karen Adelman. Pausing of RNA Polymerase II Disrupts DNA-Specified Nucleosome Organization to Enable Precise Gene Regulation. Cell, 143(4):540–551, November 2010. ISSN 00928674. doi: 10.1016/j.cell.2010.10.004.

